# Penalized generalized estimating equations for relative risk regression with applications to brain lesion data

**DOI:** 10.1101/2021.11.01.466751

**Authors:** Petya Kindalova, Michele Veldsman, Thomas E. Nichols, Ioannis Kosmidis

## Abstract

Motivated by a brain lesion application, we introduce penalized generalized estimating equations for relative risk regression for modelling correlated binary data. Brain lesions can have varying incidence across the brain and result in both rare and high incidence outcomes. As a result, odds ratios estimated from generalized estimating equations with logistic regression structures are not necessarily directly interpretable as relative risks. On the other hand, use of log-link regression structures with the binomial variance function may lead to estimation instabilities when event probabilities are close to 1. To circumvent such issues, we use generalized estimating equations with log-link regression structures with identity variance function and unknown dispersion parameter. Even in this setting, parameter estimates can be infinite, which we address by penalizing the generalized estimating functions with the gradient of the Jeffreys prior.

Our findings from extensive simulation studies show significant improvement over the standard log-link generalized estimating equations by providing finite estimates and achieving convergence when boundary estimates occur. The real data application on UK Biobank brain lesion maps further reveals the instabilities of the standard log-link generalized estimating equations for a large-scale data set and demonstrates the clear interpretation of relative risk in clinical applications.

## 1. Introduction

Often data exhibit a natural clustering, such as repeated measurements taken on the same individual over time, where the individual is considered as a cluster. The fact that the measured outcomes could be correlated within each cluster brings modelling challenges which do not arise when modelling cross-sectional data, where outcomes are typically assumed to be independent.

We are motivated by a population level imaging data set from the UK Biobank (Miller et al., 2016). Data on two brain MRI scans per individual (about 2 years apart) along with demographic and lifestyle data allow us to investigate the effect of ageing and cerebrovascular risk on the spatial distribution of brain lesions. White matter lesions (Wardlaw et al., 2013) are a common finding on MRI in older populations, but their presence is not fully explained by ageing. Lesions are associated with increased risk of stroke and dementia (Wardlaw et al., 2015) and their spatial distribution is shown to vary with cerebrovascular risk factors (Veldsman et al., 2020) in a cross-sectional analysis. When modelling such binary brain lesion maps, we should account for the potential correlation between visits within each subject, but we should also ensure the desired interpretability of the estimated regression coefficients is achieved. Here, as in many medical applications, we would like to model and interpret relative risks, i.e. a ratio of probabilities. Hence, relative risk regression would be a better choice than the widely used logistic regression, which provides log-odds ratio estimates. For example, a relative risk of 1.1 for the age effect in a particular brain region suggests that lesions in that region are 10% more likely to occur if a participant is 1 year older. Lesions are rare at the population level with most voxels (volumetric pixels) having lesion incidence below 10%, which should be accounted for in the modelling to ensure stable estimates. The modelling is to be performed at the voxel level, i.e. dependence of nearby voxels is not modelled explicitly, which is known as a mass-univariate approach. Thus, the estimated relative risks could be obtained across the brain and presented in 3D spatial maps, which allows us to explore how different risk factors impact the spatial distribution of lesions. Beyond that binary brain lesion maps application motivates the current work, the methodology we develop is generally applicable for relative risk regression in longitudinal settings.

The main approaches used for modelling longitudinal data are marginal models (also known as population average models) and conditional models (such as mixed models). The former typically uses generalized estimating equations (Liang and Zeger, 1986), and the latter maximum likelihood estimation (Laird and Ware, 1982). Detailed overviews (Fitzmaurice et al., 2011; Gardiner et al., 2009; Heagerty and Zeger, 2000) and critiques (Hubbard et al., 2010; Lee and Nelder, 2004; Lindsey and Lambert, 1998) of these methods have been made, but here we focus on the two main areas where the two approaches differ; the interpretation of the parameters and the modelling assumptions.

A mixed effects model accounts for the unobserved cluster heterogeneity by the inclusion of random effects. The assumption is that the cluster-specific parameters are random variables that are distributed according to a distribution with a few unknown fixed parameters. Thus, inference is for the population, not only for the specific cohort, and we can also obtain subject-specific effects. The assumptions of the random effects model could be inadequate, for example (i) the misspecification of the random effect distribution can influence the power of the tests or can inflate Type I error rates (Litière et al., 2007), and (ii) the random effects are typically assumed to be uncorrelated with the explanatory variables, which implies that any omitted variables are uncorrelated with the explanatory variables. As this latter assumptions cannot be validated, it has led to some authors stridently arguing against the use of mixed effects models (Hubbard et al., 2010).

In contrast, generalized estimating equations (GEEs) (Liang and Zeger, 1986) model the mean unconditionally, which implies that inference can only be made about the average effect across all subjects in the given cohort. An advantage of the GEE approach is that it does not require distributional assumptions and valid inference relies solely on the correct specification of the outcome’s mean and variance. The correlation within clusters is accounted for through the inclusion of a working correlation matrix which is treated as nuisance. The estimates of the regression coefficients are consistent, asymptotically unbiased, and asymptotically Normal when the number of clusters is sufficiently large even if the within subject correlation structure is misspecified (Liang and Zeger, 1986). However, GEE performance can suffer from small-sample bias (Sharples and Breslow, 1992; Sherman and Cessie, 1997), though bias corrections have been recently suggested by Paul and Zhang (2014). Another potential problem, discussed thoroughly for GLMs for cross-sectional binary data (Mansournia et al., 2018), is called data separation (Albert and Anderson, 1984) and it is encountered when the covariates perfectly predict the outcome. A solution to this problem for the logit link GEE is proposed by Mondol and Rahman (2019), where motivated by Firth (1993), a Jeffreys-prior penalty is used to to ensure finite estimates.

When modelling binary data, either cross-sectional, or longitudinal, a logit link function is typically selected. In medical applications, the interest often lies in interpreting relative risks as opposed to odds ratios or absolute risk differences, which naturally leads to the usage of log link. Knol et al. (2012) discussed the problem of interpreting odds ratios as relative risks in cohort studies and randomized control trials (which could be acceptable in case-control studies due to the rare disease assumption (Greenland and Thomas, 1982; Greenland et al., 1986; Knol et al., 2008), typically if the outcome incidence is less than 10%). However, it is known that the odds ratio overestimates the risk ratio (when the risk ratio is higher than 1) and the higher the outcome incidence, the larger the overestimation (Zhang and Yu, 1998). Performing a simulation study to compare 8 alternative methods for obtaining adjusted risk ratios, Knol et al. (2012) recommend the use of log-Binomial regression when adjusting for multiple covariates, and Poisson regression with robust standard errors if the log-Binomial regression fails to converge, which the authors suggest happens when the outcome incidence is high. Such convergence issues happen when the success probabilities are close to one, which leads to numerical problems in iterative estimation algorithms. A fix proposed by Carter et al. (2005) is to use a quasi-likelihood with log link, identity variance function and known dispersion for Bernoulli outcomes instead of the likelihood equations. The authors show that the resulting estimates are consistent and asymptotically Normal; similar ideas are suggested by McNutt et al. (2003) and Zou (2004). Another approach to prevent the convergence problems when using log link is proposed by Fitzmaurice et al. (2014), where a Maclaurin series approximation to the Bernoulli weights in the likelihood equations is introduced.

To ensure convergence when outcome incidence goes to 1, similar developments follow for GEE, where Zou and Donner (2013) suggest the use of the so-called ‘modified Poisson’ GEE, i.e. GEE based on the first two moments of a Poisson model using sandwich variance-covariance and log link. Even though the simulation study undertaken by Yelland et al. (2011) suggests superior convergence performance of the modified Poisson GEE when compared to GEE based on the first two moments of a Binomial model, the authors state *“Surprisingly, modified Poisson regression also failed to converge on rare occasions”*. Pedroza and Truong (2017) have also compared GEE approaches to model the relative risk and they have also reported convergence problems for the modified Poisson model for small sample size settings. We believe some of the convergence issues reported for the modified Poisson GEE could be due to data separation, which is more likely to happen for rare events and thus is not reported by Carter et al. (2005) who focus on high incidence events.

In this work we propose a practical solution for modelling repeated measures binary data with a loglink GEE. To ensure stability of the estimates for high incidence events, we base our GEE on the first two moments of a Poisson model as Carter et al. (2005) do for independent data and Zou and Donner (2013) for panel data, while allowing for the estimation of the dispersion parameter. Additionally, in previous cross-sectional analysis of binary brain lesion maps (Veldsman et al., 2020), we have observed that boundary estimates occur quite often, especially for binary covariates such as sex. Thus, to avoid the problem of boundary estimates when dealing with rare outcomes, we add a Jeffreys-prior penalty as Mondol and Rahman (2019) do for logit-link GEEs. Modelling brain lesion data implies dealing with either rare events in outer white matter, or high incidence events in the periventricular areas, so our proposed modelling approach addresses both potential convergence issues and ensures the desired risk ratio interpretability through the choice of the log link function. Note that the mass-univariate approach we take, i.e. we fit a marginal model using GEE at each voxel independently, results in 19,801 regressions across the brain in the UK Biobank application considered, highlighting the need of scalable and stable estimation methods.

In Section 2 we start by providing an overview of the standard GEE and we then describe in detail the penalized version we propose. Those two modelling approaches for relative risk regression are then applied to simulated data sets and their performance is evaluated in terms of frequency of separation occurrence as well as estimator accuracy metrics, such as bias and mean-squared error (Section 3). To demonstrate the instabilities of the vanilla GEE approach and to reveal the lucid interpretation of relative risk in medical applications, we apply the methods to a subset of the UK Biobank data, where we estimate the effect of ageing and cerebrovascular risk on lesion probability (Section 4).

## 2. Methods

First, we review the standard GEE approach and then describe the penalized version we propose to deal with boundary estimates in relative risk regression. We further outline the steps of the iterative estimation procedure and how we detect boundary estimates.

### 2.1. Generalized estimating equations

Suppose that there are *N* individuals and that each subject *i* (*i* = 1, …, *N*) has a vector of correlated binary responses ***y***_*i*_ = (*y*_*i*1_, …, *y*_*iT*_)^⊤^ at *T* time points (*t* = 1, …, *T*). Note that nothing in the present work depends on balanced data, but for notational simplicity we will assume all subjects have *T* observations. The generalized estimating equations approach introduced by Liang and Zeger (1986) explicitly accounts for the correlation between repeated measures through the specification

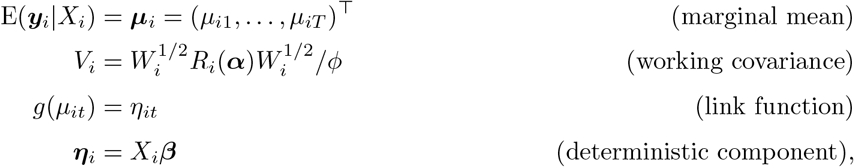

where

- *y*_*it*_ are assumed to be correlated within subject and to be independent between subjects, thus the working covariance matrix *V* is an *NT* × *NT* block-diagonal matrix with *N* blocks *V*_1_, …, *V*_*N*_.
- *R*_*i*_(***α***) is the working correlation matrix for subject *i* parameterised by parameters ***α***, *W*_*i*_ = diag{*v*_*it*_, …, *v*_*iT*_} is a *T* × *T* diagonal matrix with *v*_*it*_ = Var(*µ*_*it*_) a known variance function, and *ϕ* is the dispersion parameter, which allows for the shrinkage or inflation of the contribution of the mean to the response variance. Here we assume that *R*_*i*_(***α***) = *R*(***α***) and we subsequently suppress the index *i*.
- *g*(.) denotes the link function, which is a monotonic function that relates the marginal mean for subject *i* at time point *t* to the linear predictor *η*_*it*_.
- *X*_*i*_ denotes the *T* × *P* matrix of subject-specific covariates for subject *i*, where *X*_*i*_ collects time-specific covariate vectors ***x***_*i*1_, …, ***x***_*iT*_ in its rows and has columns ***X***_*i*1_, …, ***X***_*iP*_.
- ***β*** = (*β*_1_, …, *β*_*P*_)^⊤^ is a *P* -vector of parameters (the regression coefficients we are interested in estimating).

The generalized estimating equations, as defined by Liang and Zeger (1986), are then

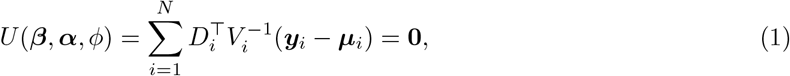

where *D*_*i*_ = *∂****µ***_*i*_*/∂****β*** is a *T* × *P* matrix and *U* is a *P* -vector. The estimation steps for the regression coefficients ***β*** as well as the ancillary association parameters ***α*** and the dispersion parameter *ϕ* are described in Section 2.3.

#### Choice of working correlation

The working correlation matrix *R*(*α*) sets the within-cluster correlation structure and it is fully characterised by the unknown parameter vector *α*. Even though its correct specification does not impact the consistency of the estimates of *β* (Liang and Zeger, 1986) and we could simply set *R* to the identity matrix, there is gain in efficiency in the estimation of *β* if the chosen dependence structure is close to the real one (Wang and Carey, 2003).

The most commonly used correlation matrices rely on a single association parameter *α*. Characteristic examples are (i) exchangeable correlation, also known as compound symmetry, where the observations within cluster share a common correlation *α*, and (ii) autoregressive correlation, where corr(*y*_*it*_, *y*_*it*_*′*) = *α*^|*t*−*t*^*′*| (*t* = 1, …, *T, t*′ = 1, …, *T*), which implies that a natural ordering of the observations exists and the correlation decays to zero as the time separation between *t* and *t*′ increases. There are other correlation structures such as Toeplitz, unstructured, etc. Crucially the choice of the correlation matrix should be considered as part of the model selection, since assuming independence can lead to quite substantial losses of efficiency (Fitzmaurice, 1995). For informative discussions on the choice of the correlation matrix, we refer the reader to Ziegler and Vens (2010) and Westgate and Burchett (2017).

### 2.2. Penalized GEE

When dealing with rare responses or small sample size, data separation is likely to occur in logistic regression (Albert and Anderson, 1984), leading to infinite maximum likelihood estimates. To ensure finiteness of the MLE in the GLM framework, adjustments to the score equations were first introduced by Firth (1993) and the finiteness of the resulting estimates in logistic regression was later proved in Kosmidis and Firth (2021). In a similar manner, Jeffreys-prior penalty, also referred to as Firth-penalty, has been proposed in the GEE framework by Mondol and Rahman (2019) as a remedy for separation in GEEs with logit link. We refer to the Mondol and Rahman (2019) method as odds ratio penalized GEE (OR-PGEE). Adding a penalty to the standard GEE in (1), the PGEE for the *p*-th regression coefficient is of the form

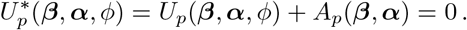

A Jeffreys-prior penalty then implies adjusting the estimating equations by a vector *A* with *p*-th component

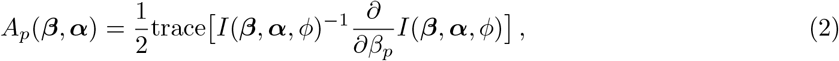

where *I*(***β, α***, *ϕ*) = E(−*∂U* (***β, α***, *ϕ*)*/∂****β***).

The motivation of our work is to ensure the direct interpretability of the estimated regression coefficients as relative risks through the usage of relative risk regression, so we propose the use of log-link function. To deal with the potential instability of the estimation algorithm for high incidence outcomes, we use a GEE with log-link and identity variance function (*v*_*it*_ = *µ*_*it*_) as Carter et al. (2005) do in cross-sectional settings, but with unknown dispersion. Because *D*_*i*_ = *W*_*i*_*X*_*i*_, the standard GEE in (1) simplifies to

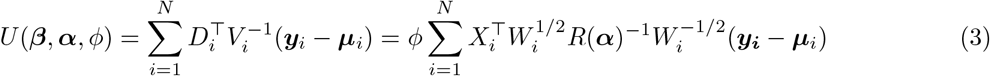

where *W*_*i*_ = diag{*µ*_*i*1_, …, *µ*_*iT*_}. We refer to the above GEE with log-link as RR-GEE.

Finally, we add a Jeffreys-prior penalty to (3) to avoid boundary estimates. Briefly, we first show that the expected negative Jacobian matrix of *U* is

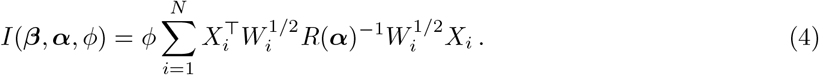

Then by further taking the derivative of *I* and substituting in expression (2), the penalty term is shown to be

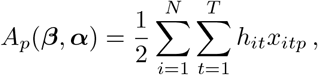

where *h*_*it*_ is the *t*-th diagonal element of the *i*-th block of the projection matrix

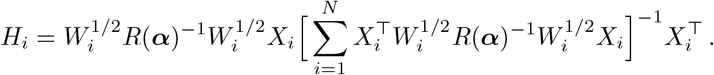

So, the modified estimating equations can be written as

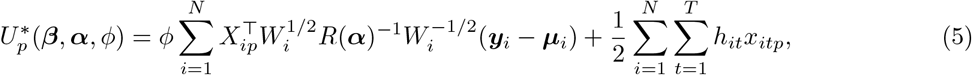

where *X*_*ip*_, ***y***_*i*_, ***µ***_*i*_ are *T* -column vectors. We refer to the above penalized estimating questions as relative risk penalized GEE (RR-PGEE). The detailed steps to obtain the form of the Jeffreys-prior penalty term *A*_*p*_(***β, α***) for this RR-GEE are included in Appendix A.

### 2.3. Parameter estimation

In a similar manner to maximum likelihood parameter estimation in GLMs, we adopt an iterative procedure to solve the estimating equations (5) for RR-PGEE. The iterative process is as suggested by Liang and Zeger (1986) and it entails repeated use of a quasi Newton-Raphson iteration for the estimation of ***β*** and method of moments estimation of ***α*** and *ϕ*. The resulting estimates 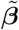 of ***β*** are the RR-PGEE estimates along with 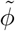 and 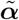. The iterative procedure is as follows

*Step 1* Initialization.

Set the intercept term to the log mean response incidence 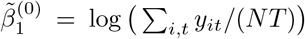 and 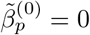 for *p* = 2, …, *P*.

*Step 2* For *k* = 1, …, *K*, repeat the following two steps until a convergence criterion is satisfied, e.g. 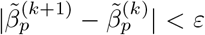 for all parameters *β*_*p*_, *p* = 1, …, *P* and positive convergence tolerance *ε*, or the maximum number of iterations *K* has been reached without convergence.

i. Given 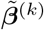 and assuming exchangeable working correlation, i.e. all off-diagonal elements of *R*(***α***) are equal to *α*:

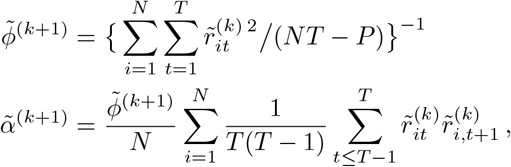

where the Pearson residuals at iteration *k* are 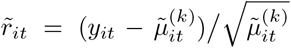 and 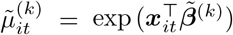.
ii. Using 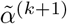 and 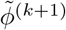, the quasi Newton-Raphson update is

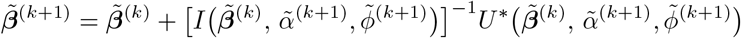

*Step 3* Set 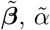 and 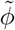 to the values at the final iteration of Step 2.

Using the same steps as above, we can also obtain the RR-GEE estimates 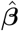, where *U* as in Equation (3) is used in the Newton-Raphson update in Step 2(ii) instead of *U* ^∗^. Both sets of estimates 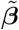 and 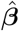 are obtained using the log-link function and identity variance function with unknown dispersion.

#### Robust variance

To estimate the variance of the estimated regression coefficients 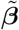, the sandwich variance-covariance matrix 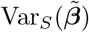 proposed by Liang and Zeger (1986), also called robust variance, is used

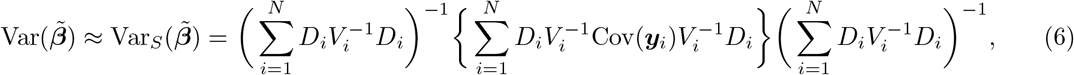

where to obtain the variance estimate 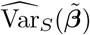, we replace Cov(***y***_*i*_) by (***y***_*i*_ − ***µ***_*i*_)(***y***_*i*_ − ***µ***_*i*_)^⊤^ and ***β***, *ϕ* and *α* by their estimates; the same is done to obtain 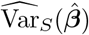. The consistency of 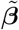 and 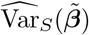 does not depend on the correct choice of the working correlation matrix *R*, but only on the correct specification of the outcome mean regression (Liang and Zeger, 1986).

#### Detection of boundary estimates

To detect boundary estimates, we define the boundary estimates criterion (BEC) for the *p*-th regression coefficient at iteration *k*

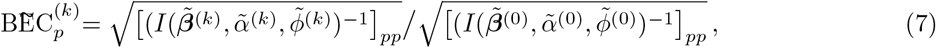

where *k* = 1, …, *K* and *I* is the expected negative Jacobian matrix as defined in Equation (4). Similarly, we define BÊC by replacing 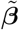 with 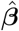 as well as the estimates of *α* and *ϕ*.

Boundary estimates occur if this ratio diverges as *k* grows for any of the *P* estimated parameters. The ratio depends on the expected negative Jacobian of the estimating equations as given in Equation (4), which gives us information on the curvature of the estimating function *U*. To give some intuition behind this criterion, in the single parameter setting, when boundary estimates occur for large values of the parameter *β* the Jacobian will have value almost zero, thus *I*^−1^ will be diverging to large values. For *k* = 1, …, *K*, we monitor 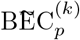, and if it diverges as *k* grows for any of the *P* estimated parameters, it suggests we have detected boundary estimates; similarly for 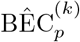.

## 3. Simulation study

To compare the performance of our proposed penalized GEE for relative risk regression (RR-PGEE) and the associated standard GEE (RR-GEE), we perform a simulation study in a similar manner to Mondol and Rahman (2019). The main aim of the simulation study is to investigate the frequency of boundary estimates for correlated binary data and to evaluate how the modified GEE performs in comparison to the standard GEE in those cases.

### 3.1. Simulation setup

We simulate balanced correlated binary data *y*_*it*_ for subject *i* at time point *t* (*i* = 1, …, *N, t* = 1, …, *T*). Specifying our model as per Section 2.1, the deterministic component of our generative model is of the form

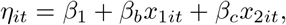

where

- *β*_1_ is the intercept term, which determines the response incidence in the data set.
- a binary covariate *X*_1_ is considered, which is time-invariant (e.g. sex) and its values are sampled from Bernoulli distribution with probability of success *c*. The inclusion of a time-invariant binary covariate could potentially lead to separation and the regression coefficient *β*_*b*_ and its standard error could be diverging to infinity.
- a continuous covariate *X*_2_ is considered, which is time-varying with equally-spaced values {0.2, 0.4, 0.6, …} for each subject *i*.

As the link function is log-link, the marginal mean is *µ*_*it*_ = exp (*η*_*it*_). By specifying the marginal mean and variance of ***y*** and the correlation matrix *R*(***α***), the method proposed by Qaqish (2003) is used to simulate correlated binary data. Briefly, Qaqish introduced a family of multivariate Bernoulli distributions with a conditional linear property, which provides an efficient approach to simulate correlated binary variables.

The code available as part of the `binarySimCLF R` package has been adjusted to account for our choices of link function and identity variance function Var(*µ*_*it*_) = *µ*_*it*_.

The fixed simulation parameters are as follows (i) fixed number of time points *T* = 4, resulting in ***x***_2*i*_ = (0.2, 0.4, 0.6, 0.8)^⊤^ across all subjects *i*, with the regression coefficient *β*_*c*_ = 0.2; (ii) correlation structure is set to exchangeable with correlation parameter *α*. The base simulation has parameters fixed at: regression coefficients ***β*** = (*β*_1_, *β*_*b*_, *β*_*c*_)^⊤^ = (−4, 1.6, 0.2)^⊤^, sample size *N* = 50, time-invariant binary covariate proportion of 1’s set to *c* = 0.2, within-subject correlation *α* = 0.4. We vary each of five simulation parameters one at a time, keeping the others fixed as above, as follows

- *β*_1_ ∈ {−4, −3, −2}: the overall response incidence, with smallest value giving rare events and greatest chance of boundary estimates,
- *β*_*b*_ ∈ {1.2, 1.4, 1.6, 1.8, 2.0}: the binary covariate effect, with smaller effects increasing the chance of boundary estimates,
- *c* ∈ {0.2, 0.3, 0.4, 0.5, 0.6, 0.7, 0.8}: the proportion of 1’s for the binary time-invariant covariate, with higher proportions increasing the risk of boundary estimates if the response incidence is low,
- *α* ∈ {0.2, 0.3, 0.4, 0.5, 0.6, 0.7, 0.8}: the strength of the within-subject correlation,
- *N* ∈ {25, 50, 75, 100, 500, 1000}: the sample size, with the smallest sample sizes resulting in higher chance of boundary estimates.

### 3.2. Model evaluation

We simulate *R* = 1, 000 data sets for each of the simulation scenarios listed above. For each repetition *r* (*r* = 1, …, *R*), RR-GEE and RR-PGEE are fitted to obtain estimates 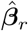 and 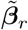, respectively, along with their sandwich variance and boundary estimates criterion values. Note that for the simulation study, the true value of the dispersion parameter *ϕ* is set to 1, but it is estimated using RR-GEE or RR-PGEE when fitting the marginal models. The tolerance for convergence is set to *ε* = 10^−4^ and the maximum number of iterations to *K* = 25.

To judge whether boundary estimates occur, we focus on the boundary estimates criterion (BEC) in Equation (7) obtained from fitting RR-GEE and we keep record of those values at each iteration of the IRLS algorithm. We state that boundary estimates occurred if BEC at the final iteration is greater than 10 for any of the three regression coefficients. Note that for some of the simulated data sets, either of the modeling approaches could return missing values due to numerical instabilities (referred to as ‘failed IRLS’), or it can fail converging for the maximum number of permitted iterations *K* (referred to as ‘non-converging’).

To compare the two modelling approaches in terms of estimator accuracy, the bias and mean-squared error (MSE) of the regression coefficient *β*_*b*_ are estimated. For RR-GEE, bias is calculated as 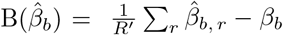 and MSE as 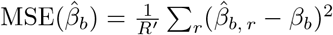, where 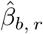 is the RR-GEE estimate for the *r*-th simulated data set, *β*_*b*_ is the true value, and summation is over the *R*′ data sets where estimation did not fail. Similarly, 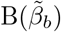 and 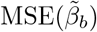 are obtained using RR-PGEE estimates 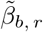 across repetitions. To further explore the performance of the variance estimators of the regression coefficients, we compare the average sandwich variance 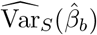 with the variance of the estimates 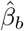 across repetitions for RR-GEE and similarly for RR-PGEE 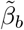. If the sandwich variance correctly estimates the variance of the estimators, the ratio of those two quantities should be close to 1.

### 3.3. Results

#### Motivating example

As an illustrative example, we show the results from fitting RR-GEE and RR-PGEE to the 1,000 simulated data sets under the base simulation setup, i.e. ***β*** = (*β*_1_, *β*_*b*_, *β*_*c*_)^⊤^ = (−4, 1.6, 0.2)^⊤^, *N* = 50, *c* = 0.2, *α* = 0.4.

On 233 out of 1,000 simulated data sets boundary estimates occurred, meaning at least one of the three regression coefficients has BEC greater than 10. Note that the threshold of 10 is chosen empirically, for a histogram of the BEC values for *β*_*b*_, see Figure B.1. We randomly selected one of the 233 boundary estimates data sets and the results are summarized in Table 1. Increasing the number of iterations highlights the rapid increase in BEC as well as the elevated regression coefficient and z-score for the binary covariate obtained using RR-GEE, where 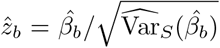, while the estimates based on RR-PGEE are more stable and the iterative algorithm converges after 10 iterations. One warning here is that if we explore the *z*-scores for the data sets where boundary estimates occur, we get large absolute values of 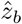 for most of those data sets when fitting RR-GEE. The reality is that users tend to extract the *z*-scores without realizing that software packages handle boundary estimates differently and that the user may receive no warning about boundary estimates.

**Table 1:**
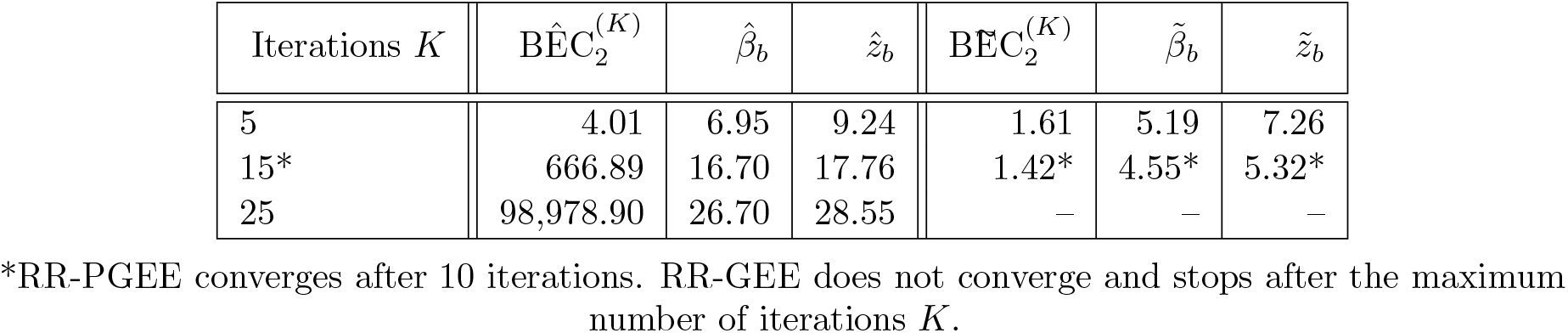
Performance of one boundary estimates data set for the effect of the binary covariate (*p* = 2) across *K*={5, 15, 25} maximum number of iterations. The simulation parameters are set to ***β*** = (*β*_0_, *β*_*b*_, *β*_*c*_)^⊤^ = (−4, 1.6, 0.2)^⊤^, *N* = 50, *c* = 0.2, *α* = 0.4.

Of the remaining 767 data sets, 18 had numerical underflow or non-convergence failure for either of the methods. Figure 1 explores the remaining 749 estimated coefficients for converging data sets with finite estimates. The highest density of points is close to the true value of *β*_*b*_ = 1.6 for both RR-GEE and RR-PGEE, with RR-PGEE demonstrating shrinkage to the true value. Also, RR-PGEE leads to slightly higher *z*-scores than RR-GEE due to the shrinkage of the standard errors towards zero.

**Figure 1:**
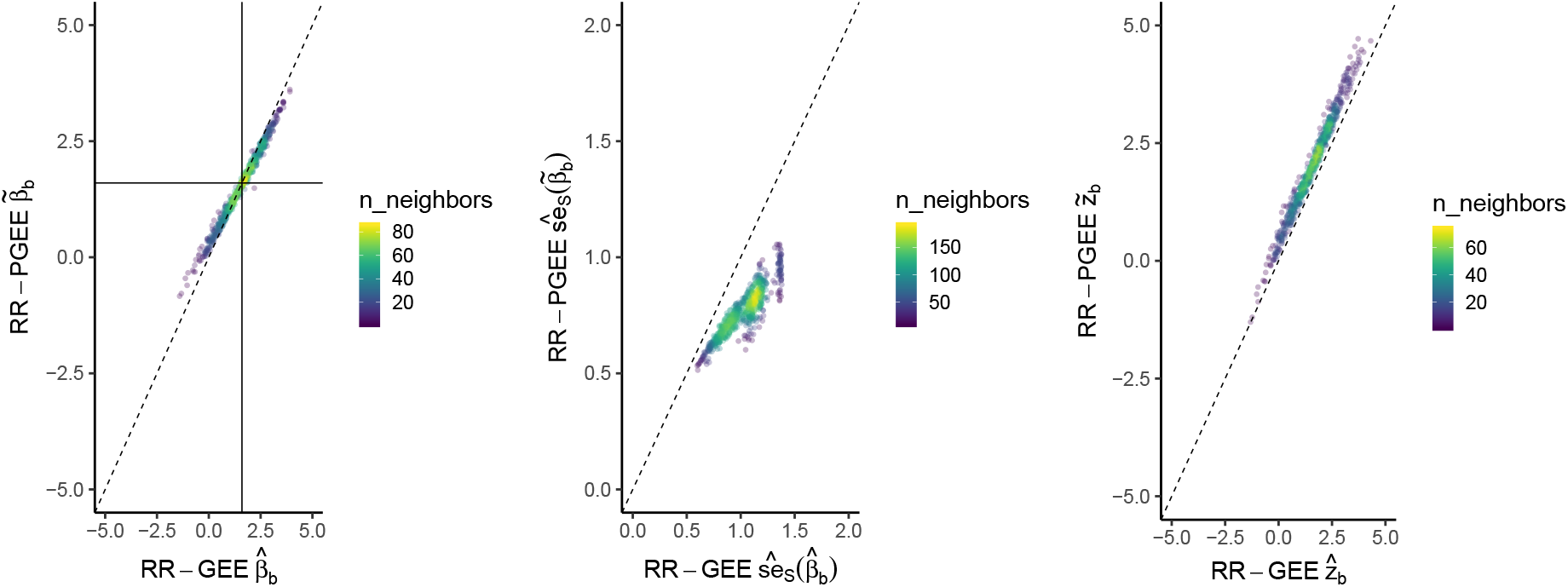
Illustration of converging data sets with finite estimates. (Left) Estimated regression coefficients for the binary covariate *β*_*b*_ estimated from 749 data sets, with solid lines at the true value of 1.6, (Middle) sandwich standard error, and (Right) estimated *z*-scores (estimated coefficient divided by sandwich standard error) for RR-PGEE vs RR-GEE. Dashed line is the identity line; those are density plots, i.e. the brighter the colour, the higher the density of the points. The plot summarises the 749 non-separated data sets out of 1,000 data sets simulated under the base simulation scenario: ***β*** = (*β*_1_, *β*_*b*_, *β*_*c*_)^⊤^ = (−4, 1.6, 0.2)^⊤^, *N* = 50, *c* = 0.2, *α* = 0.4.

#### Boundary estimates

The convergence issues we run into when fitting both RR-GEE and RR-PGEE are summarized in Table 2. The number of boundary estimates data sets out of 1,000 simulated under each scenario increases: (i) the rarer the event as controlled by the intercept term *β*_1_, (ii) the weaker the effect of the binary covariate *β*_*b*_, (iii) the stronger the association parameter *α*, (iv) the higher the proportion of 1’s for the binary covariate, (v) the lower the sample size *N*. Given that the main aim of the penalized GEE for relative risk regression is to ensure finite estimates, our results reveal that the BEC value is above the chosen threshold of 10 for a few of the data sets but IRLS always converges. It happens that IRLS fails for RR-PGEE as it does for RR-GEE, but it happens less frequently for most simulation scenarios. Overall, RR-PGEE performs well when boundary estimates are detected, i.e. when the BEC value for RR-GEE diverges. Note that for some data sets, both methods do not converge for the specified maximum number of iterations and the frequency of this happening is similar between the methods; a further check into those data sets (as in the last column of Table 2) shows that the outcome incidence is exactly zero for most of those data sets.

**Table 2:**
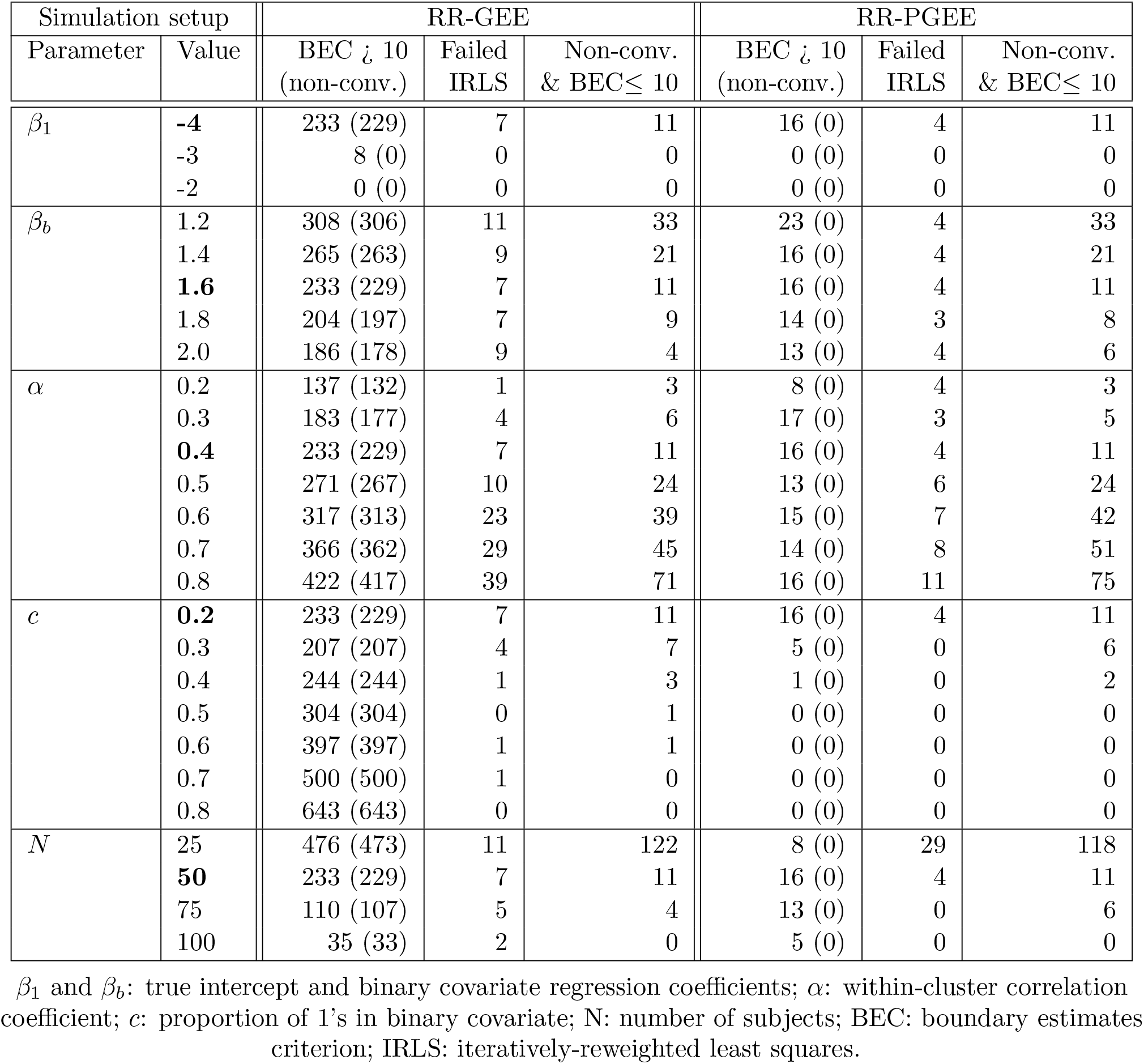
Simulation performance across simulation scenarios. The simulation parameters are set to ***β*** = (*β*_1_, *β*_*b*_, *β*_*c*_)^⊤^ = (−4, 1.6, 0.2)^⊤^, *N* = 50, *c* = 0.2, *α* = 0.4 (in bold) and each row of the table summarizes 1,000 replications with just one parameter altered, e.g. the first block of rows represents scenarios when *β*_1_ is altered and all other parameters stay fixed.

#### Estimator accuracy

In Table 3, we present the bias and mean-squared error of the binary covariate estimates 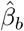 and 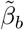. We explore the data sets with finite and converging estimates to investigate how the penalty impacts the estimates, but we also look at the RR-PGEE unconditional summaries including estimates for boundary estimates data sets (in parenthesis). The results reveal that the bias is comparable between the two modelling approaches and RR-PGEE has lower MSE across the simulation scenarios, except for high proportion of 1’s for the binary covariate (*c* ∈ {0.6, 0.7, 0.8}, Table 3). The response incidence is very low (exp(*β*_1_) = exp(−4) = 0.02), so if we further explore sample sizes *N* = 100 and *N* = 500 for varying proportions of 1’s in the binary covariate (see Table B.1), the MSE and bias performance is encouraging, i.e. both are decreasing when sample size increases indicating consistency of the RR-PGEE estimates.

**Table 3:** Bias and mean-squared error of the binary covariate coefficient *β*_*b*_ across simulation scenarios. The simulation parameters are set to ***β*** = (*β*_1_, *β*_*b*_, *β*_*c*_)^⊤^ = (−4, 1.6, 0.2)^⊤^, *N* = 50, *c* = 0.2, *α* = 0.4 (in bold) and each row of the table summarizes 1,000 replications with just one parameter altered, e.g. the first block of rows represent scenarios when *β*_1_ is altered and all other parameters stay fixed. Summaries included are conditional on both estimation methods converging and the estimates being finite (no boundary estimates), with unconditional summaries for RR-PGEE included in parenthesis.

| Simulation setup |  | Finite & converging |  |  |  |  |
| --- | --- | --- | --- | --- | --- | --- |
| Parameter | Value | N.sim | B($\hat{\beta}_b$) | B($\tilde{\beta}_b$) | MSE($\hat{\beta}_b$) | MSE($\tilde{\beta}_b$) |
| $\beta_1$ | **-4** | 749 (969) | -0.04 | -0.05 (0.19) | 0.88 | 0.61 (2.18) |
|  | -3 | 992 (1000) | 0.03 | 0.03 (.04) | 0.53 | 0.42 (0.50) |
|  | -2 | 1000 (1000) | 0.03 | 0.02 (0.02) | 0.12 | 0.11 (0.11) |
| $\beta_b$ | 1.2 | 648 (940) | 0.12 | 0.15 (0.11) | 0.89 | 0.62 (2.23) |
|  | 1.4 | 705 (959) | 0.04 | 0.05 (0.16) | 0.87 | 0.61 (2.26) |
|  | **1.6** | 749 (969) | -0.04 | -0.05 (0.19) | 0.88 | 0.61 (2.18) |
|  | 1.8 | 780 (975) | -0.05 | -0.09 (0.23) | 0.82 | 0.58 (2.07) |
|  | 2.0 | 801 (977) | -0.09 | -0.15 (0.24) | 0.82 | 0.61 (2.02) |
| $\alpha$ | 0.2 | 859 (985) | 0.02 | $-4 \times 10^{-3}$ (0.12) | 0.74 | 0.54 (1.51) |
|  | 0.3 | 807 (975) | 0.01 | -0.01 (0.14) | 0.82 | 0.58 (1.77) |
|  | **0.4** | 749 (969) | -0.04 | -0.05 (0.19) | 0.88 | 0.61 (2.18) |
|  | 0.5 | 695 (957) | -0.04 | -0.05 (0.18) | 0.87 | 0.60 (2.36) |
|  | 0.6 | 621 (936) | -0.04 | -0.05 (0.22) | 0.90 | 0.60 (2.67) |
|  | 0.7 | 560 (927) | -0.04 | -0.06 (0.24) | 0.91 | 0.60 (2.90) |
|  | 0.8 | 468 (898) | 0.04 | -0.01 (0.29) | 0.95 | 0.60 (3.13) |
| $c$ | **0.2** | 749 (969) | -0.04 | -0.05 (0.19) | 0.88 | 0.61 (2.18) |
|  | 0.3 | 782 (989) | -0.09 | -0.18 (0.24) | 0.85 | 0.64 (1.85) |
|  | 0.4 | 752 (997) | -0.19 | -0.33 (0.20) | 0.84 | 0.69 (1.58) |
|  | 0.5 | 695 (1000) | -0.27 | -0.47 (0.14) | 0.76 | 0.72 (1.31) |
|  | 0.6 | 601 (1000) | -0.43 | -0.67 (0.02) | 0.84 | 0.90 (1.12) |
|  | 0.7 | 499 (1000) | -0.64 | -0.92 (-0.14) | 1.03 | 1.27 (0.96) |
|  | 0.8 | 357 (1000) | -0.93 | -1.26 (-0.42) | 1.46 | 1.99 (0.89) |
| $N$ | 25 | 391 (845) | -0.02 | -0.05 (0.56) | 0.84 | 0.61 (4.04) |
|  | **50** | 749 (969) | -0.04 | -0.05 (0.19) | 0.88 | 0.61 (2.18) |
|  | 75 | 881 (981) | 0.04 | 0.02 (0.14) | 0.83 | 0.60 (1.40) |
|  | 100 | 963 (995) | 0.05 | 0.03 (0.07) | 0.75 | 0.56 (0.82) |
| | 500 | 1000 (1000) | $7 \times 10^{-4}$ | -0.01 (-0.01) | 0.13 | 0.12 (0.12) |
| | 1000 | 1000 (1000) | $7 \times 10^{-3}$ | $4 \times 10^{-3}$ ($4 \times 10^{-3}$) | 0.06 | 0.06 (0.06) |

B: bias; MSE: mean squared error.

Table 4 reports on the variance properties, showing that RR-PGEE has smaller variance in all settings considered, i.e.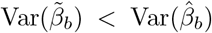. The accuracy of the estimated variance is assessed through the relative variance, i.e. mean of 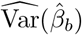 across repetitions divided by variance of the estimated coefficients 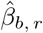(Table 4), and similarly for 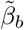. For the finite and converging data sets, the RR-GEE seems to be slightly overestimating the true variance of the binary covariate coefficient and RR-PGEE is slightly underestimating it, with ratios approaching 1 as *N* increases.

**Table 4:**
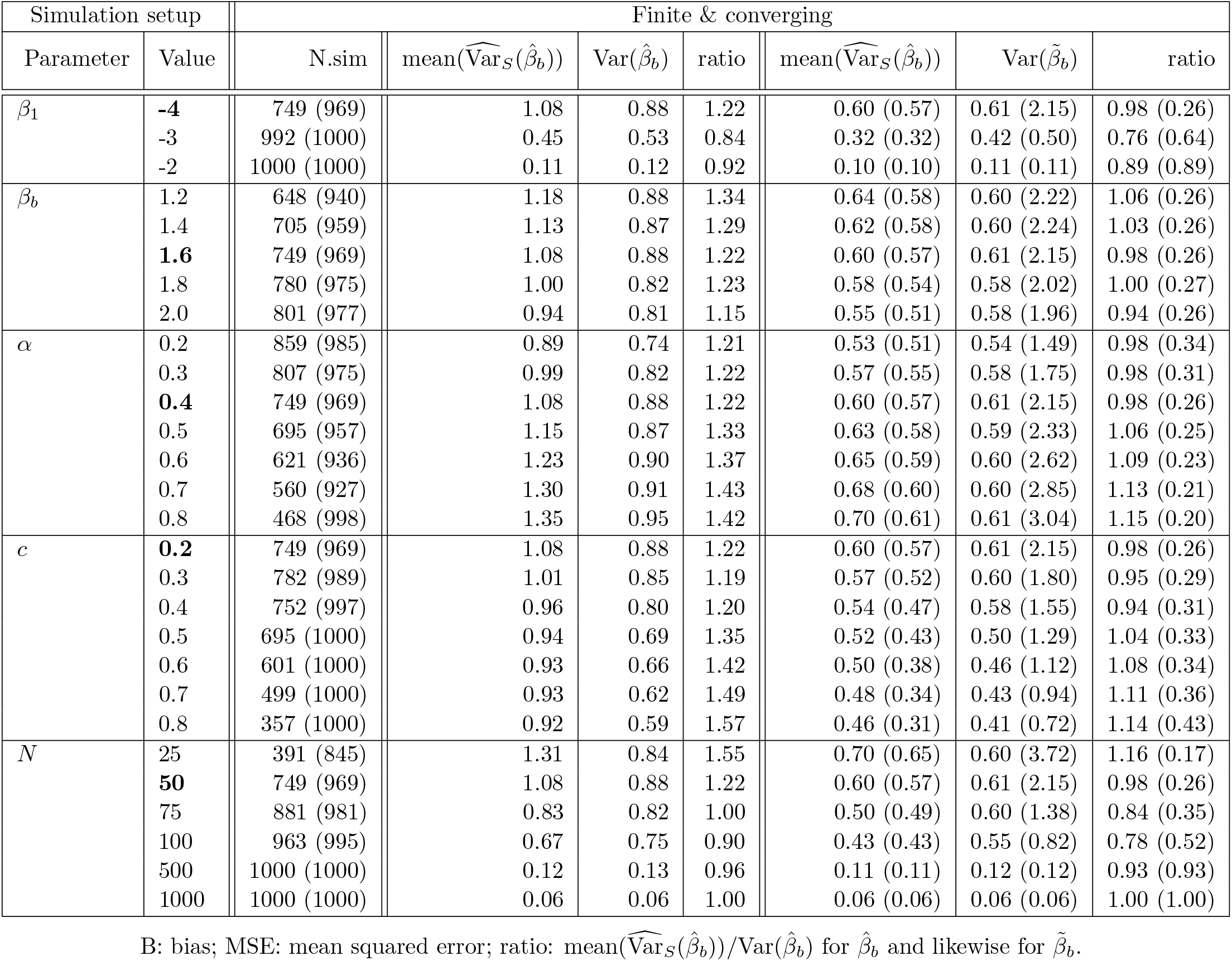
Accuracy of variance estimates of the binary covariate coefficient *β*_*b*_ across simulation scenarios. We compare the average sandwich variance 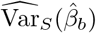 with the variance of the estimates 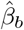 across N.sim repetitions for RR-GEE and the same for RR-PGEE, i.e.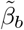. The simulation parameters are set to ***β*** = (*β*_1_, *β*_*b*_, *β*_*c*_)^⊤^ = (−4, 1.6, 0.2)^⊤^, *N* = 50, *c* = 0.2, *α* = 0.4 (in bold) and each row of the table summarizes 1,000 replications with just one parameter altered, e.g. the first block of rows represent scenarios when *β*_1_ is altered and all other parameters stay fixed. Summaries included are conditional on both estimation methods converging and the estimates being finite (no boundary estimates), with unconditional summaries for RR-PGEE included in parenthesis.

Note that the unconditional summaries for RR-PGEE (in parenthesis) exhibit stable performance for the data sets where boundary estimates are detected for RR-GEE. Furthermore, MSE and bias values decrease towards 0 and variance ratios go to 1 with increasing *N*, which we would expect from a consistent estimator.

## 4. Application to brain lesion data

### 4.1. Data

To demonstrate the performance of the two methods for relative risk regression - RR-GEE and RR-PGEE - we apply them to a subset of the UK Biobank data set (Miller et al., 2016). White matter lesions are an extremely common finding on MRI in older adults, with age being the strongest predictor of lesions, followed by cerebrovascular risk (CVR) factors. CVR factors include smoking and hypertension; below we define a CVR score using 6 variables. It is known that the presence of CVR increases the risk of stroke and dementia (Wardlaw et al., 2015). Thus, we aim to investigate the effect of ageing and cerebrovascular risk on the spatial distribution of lesions, exploring both the cross-sectional and longitudinal effects.

Participants are selected according to the flow chart in Figure 2 resulting in a data set of 1,578 healthy ageing individuals; the same selection criterion is used as in Veldsman et al. (2020). The data available for the analysis include binary brain lesion maps at two time points along with data on cerebrovascular risk factors and other variables.

**Figure 2:**
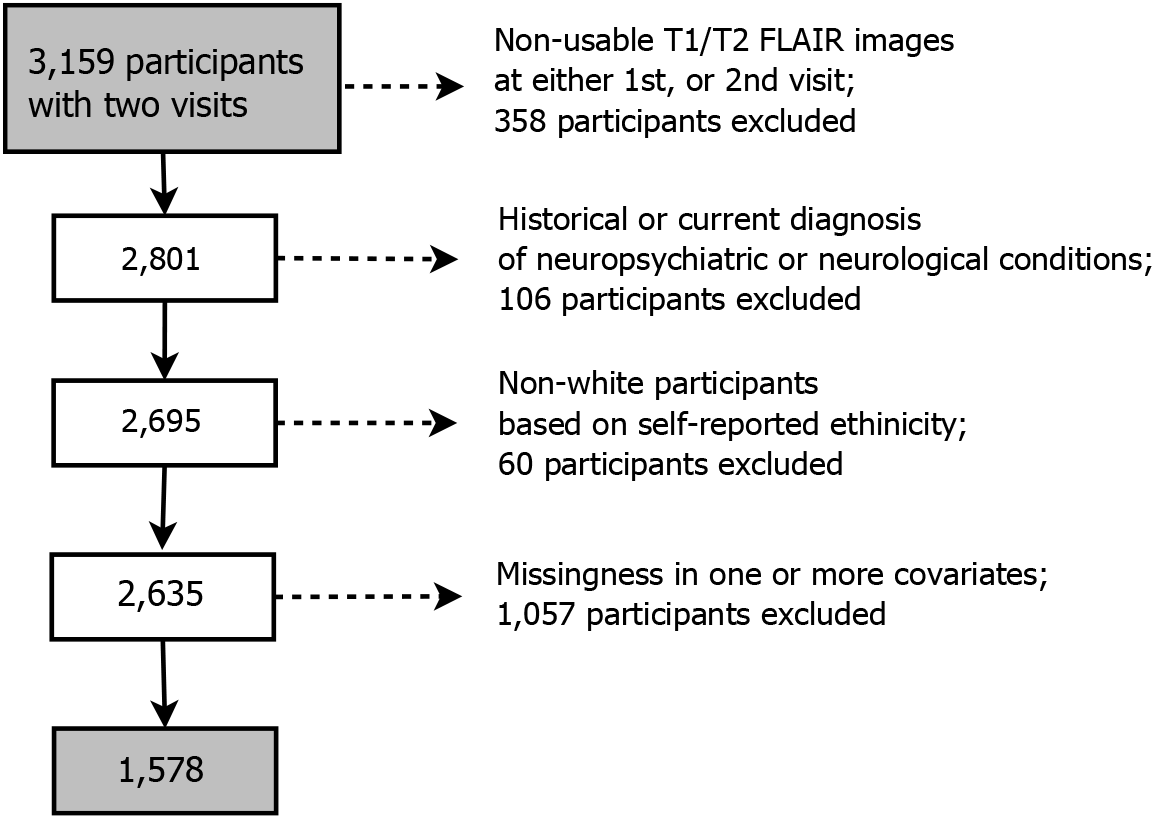
Diagram demonstrating the flow of gradually refining participants starting from all UK Biobank participants with available T1 and T2 FLAIR structural brain images at the two imaging visits. Most common neuropsychiatric/neurological conditions in decreasing number of participants: stroke (29), transient ischaemic attack (18), etc. (see Table B.2 for a full list).

#### MRI data

To generate the binary brain lesion maps, we use two MRI imaging modalities (T1-weighted and T2-weighed FLAIR) as input to the lesion segmentation software BIANCA (Griffanti et al., 2016). The resulting lesion maps are in native space (i.e. subject-specific) with 1 indicating the presence of a lesion and 0 the absence. To facilitate the analysis, those binary maps are first registered to a common space (2 × 2 × 2mm^3^ Montreal Neurological Institute (MNI) template) by applying spatial normalisation, and are then binarised with a 0.5 threshold. The image preprocessing, the lesion segmentation step as well as the derivation of the estimated spatial normalisation parameters are part of the published UKB preprocessing pipeline (Alfaro-Almagro et al., 2018). Binary lesion maps of dimension 91 × 109 × 91 voxels (volumetric pixels), are available for 1,578 subjects at two time points. The lesion volume at each time point is also available as part of the UKB variables’ catalogue.

#### Cerebrovascular risk score

A cerebrovascular risk score is calculated as a sum of six categorical variables representing six risk factors, as described by Veldsman et al. (2020). The risk factors include hypertension, hypercholesterolemia, smoking, diabetes, waist-to-hip ratio and the APOE-*ε* (apolypoprotein-E) status, with categorical variables (0/1/2) for smoking and APOE-*ε* status, and binary variables (1 indicating presence of the risk factor) for the rest of the risk factors. The resulting score can range from 0 to 8 and is computed for both time points.

#### Confounding variables

The minimal set of confounding variables, as suggested by Alfaro-Almagro et al. (2020), includes age, sex, age by sex interaction and head size scaling.

### 4.2. Analysis setup

We have correlated binary data on *N* = 1, 578 subjects at *T* = 2 time points, where each subject *i* comes with a binary map *y*_*it*_(*s*) ∈ {0, 1}, *s* ∈ ℬ ⊂ ℝ^3^ at time point *t*. ℬ is the human brain and we consider *M* cubic cells *s*_*j*_ (voxels) as a discretization of the 3D brain on a regular rectangular grid, where *s*_*j*_ denotes the *j*th voxel within the brain (*j* = 1, …, *M*). We model the available binary lesion maps voxel-wise, i.e. we fit a model at each voxel marginally, ignoring the spatial dependence in the brain, in what is known as a mass-univariate approach. Using the same notation as in Section 2, the marginal model is of the form

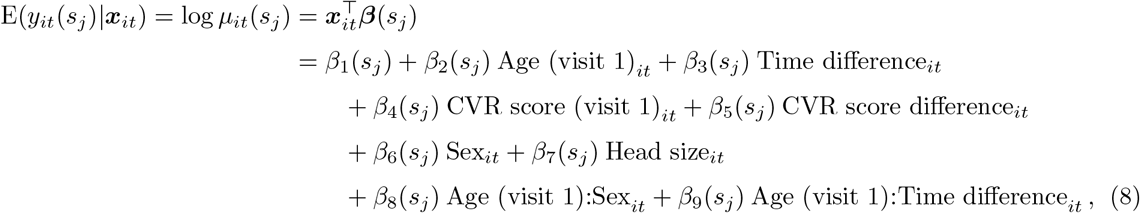

where ***x***_*it*_ is a *P* -vector of time-specific covariates for subject *i* at time *t*. The proposed model above includes eight explanatory variables and *P* = 9 with an intercept. We follow the recommended practice of splitting time-varying variables into pure cross-sectional and pure longitudinal variables (Neuhaus and Kalbfleisch, 1998). This leads to five time-invariant variables, i.e. their value is the same across the *T* = 2 time points, namely Age (visit 1), CVR score (visit 1), Sex, Head size (average across two visits) and Age (visit 1) by Sex interaction. The remaining three explanatory variables are time-varying; Time difference captures the longitudinal effect of age and is constructed as a *T* -vector (Time difference_*i*1_, Time difference_*i*2_)^⊤^ = (0, Age (visit 2)_*i*_ − Age (visit 1)_*i*_)^⊤^ for subject *i*; similarly for CVR score and Age (visit 1) by Time difference interaction. Note that prior to fitting the model using RR-GEE and RR-PGEE, Age (visit 1), CVR score (visit 1) and Head size are demeaned across all subjects.

At each voxel, we obtain RR-GEE estimates 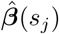 and RR-PGEE estimates 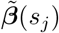 along with sandwich standard errors and the associated *z*-scores. We also keep track of the boundary estimates criterion (as defined in Equation (7)) to detect boundary estimates. To explore the interpretability of the estimated coefficients and the choice of the link function, we fit the penalized GEE for logistic regression (OR-PGEE) as introduced by Mondol and Rahman (2019) and obtain a *P* -vector of estimates ***β***^*^(*s*_*j*_) across voxels *s*_*j*_. The first two moments of the response are set to E(*y*_*it*_) = *µ*_*it*_ and Var(*y*_*it*_) = *µ*_*it*_(1 − *µ*_*it*_)*/ϕ* and the link function used is logit. We have adjusted the Mondol and Rahman (2019) `R` code to use the same starting values as ours and to use the same convergence criterion. For rare events, we would expect the estimated relative risk and odds ratios to be highly similar. We also transform the estimated log-odds to relative risks to allow for further comparisons. For example, the transformation to get the relative risk for Age at visit 1 from the estimated log-odds ***β***^*^(*s*_*j*_) is of the form 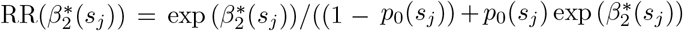, where 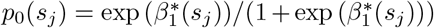 and 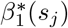 is the log-odds for the intercept term and 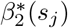 for age, respectively.

In contrast to the simulation study, we allow for the estimation of the dispersion parameter *ϕ*(*s*_*j*_) across voxels *s*_*j*_. We use exchangeable correlation matrix with parameter *α*(*s*_*j*_) to measure the strength of the within-subject correlation. The convergence tolerance for all three methods is set to *ε* = 0.001 with maximum number of iterations set to *K* = 25.

Lesions are known to be disproportionately located in periventricular or in deep white matter regions, with varying lesion incidence across the brain. This, in a similar way to cross-sectional voxel-wise analysis (Lampe et al., 2019; Rostrup et al., 2012; Veldsman et al., 2020), we exclude voxels when the lesion count is too low. We have chosen 6 as our threshold, i.e. the models are fitted only at voxels where 6 or more lesions are present across all subjects and both visits. To identify the voxels of interest, we create maps of the lesion incidence across the brain, where *p*_1_(*s*_*j*_) denotes the lesion incidence at voxel *s*_*j*_ for visit 1 and *p*_2_(*s*_*j*_) for visit 2, respectively, and ***p***_1_ and ***p***_2_ denote the corresponding images.

Given the large number of regressions performed across the brain, when exploring the *z*-scores we report results based on a fixed threshold of ±1.96 at 5% significance level, or we adopt a false discovery rate (FDR) correction (Benjamini and Hochberg, 1995) at 5% significance level. To gain better understanding of the localized effect of ageing and CVR score on lesion occurrence both cross-sectionally and longitudinally, we also explore spatial maps of the unthresholded/uncorrected (or ‘raw’) *z*-scores and relative risks, i.e. exploring all voxels where the models are fitted. Note that the chosen log link function allows us to interpret the exponent of the estimated coefficients directly as the relative risk.

### 4.3. Results

The analysis to follow is based on UKB data on 1,578 individuals (mean age 62.7 ± 7.2 years, 781 men) with sample characteristics included in Table 5. The summaries for CVR score and lesion volume across the two visits suggest a slight increase in both over time.

**Table 5:** Characteristics of UK Biobank dataset of 1,578 participants.

| Characteristics | Mean (SD) | Median (range) |
| --- | --- | --- |
| Age, visit 1 (years) | 62.7 (7.2) | 62.9 (50.0; 79.1) |
| Time between visits (years) | 2.2 (0.1) | 2.2 (2.0; 2.8) |
| Sex (baseline female) | 49.5% Men | — |
| Head size scaling* | 1.3 (0.1) | 1.3 (0.9; 1.7) |
| CVR score, visit 1 | 1.7 (1.3) | 2.0 (0; 6) |
| CVR score, visit 2 | 1.8 (1.2) | 2.0 (0; 7) |
| Lesion volume, visit 1 (mm<sup>3</sup>) | 4,626 (5,327) | 2,813 (94; 53,608) |
| Lesion volume, visit 2 (mm<sup>3</sup>) | 5,050 (5,894) | 3,024 (136; 54,713) |

\*average of head size scaling for visits 1 and 2
N: number of participants; SD: standard deviation; CVR: cerebrovascular risk.

#### Spatial distribution of lesions

The lesion incidence across 1,578 participants at their first scan (Figure 3) highlights the periventricular and deep white matter regions as expected. The empirical relative risk ***p***_2_*/****p***_1_ suggests that the lesion incidence increases from visit 1 to visit 2 for some deep white matter areas. 19,801 voxels with six or more lesions present across all subjects and both visits are considered for the analysis, i.e. 19,801 regressions across the brain as outlined in Section 4.2.

**Figure 3:**
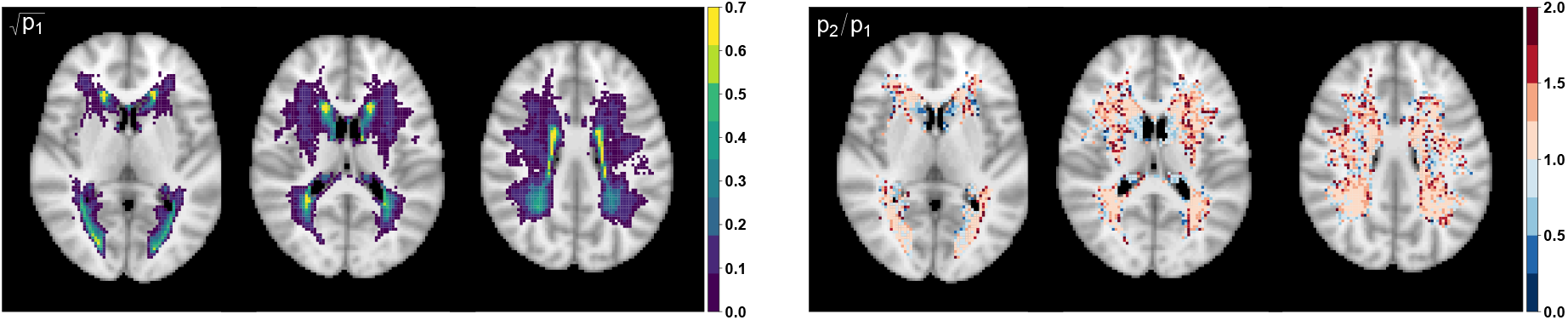
Square–root transformed empirical lesion incidence ***p***_1_ for visit 1 (left) and empirical relative risk ***p***_2_*/****p***_1_ (right) based on binary lesion maps of 1,578 UKB participants across two visits; axial slices *z* = {40, 45, 50} shown from left to right. 19,801 voxels with six or more individuals having a lesion across two visits are plotted with the remaining voxels plotted as transparent to show the standard anatomical MRI for reference. Square-root transformed incidence is used to better visualise the structure in the low incidence regions.

#### Boundary estimates

As described in Section 4.2, the selected marginal model is fitted using RR-GEE and RR-PGEE at 19,801 voxels across the brain. Boundary estimates are more likely to occur for binary covariates but we monitor the BEC values as defined in Equation (7) to detect boundary estimates for any of the covariates in the model. The threshold on BEC values is chosen empirically, see Figure B.2. So, we set the threshold for BEC to 10 as we did for the simulations in Section 3. We get diverging standard errors for 4% of the voxels (763 voxels out of 19,801) and missing values for 1% (198 voxels) for RR-GEE and 90 voxels and 2 missing values for RR-PGEE, respectively. However, the iterative algorithm converges for 350 out of those 763 voxels for RR-GEE and for 58 out of 90 for RR-PGEE, which suggests that the threshold for the boundary estimates detection criterion may be too conservative. If we increase the threshold to 100, 337 voxels have boundary estimates (2%) for RR-GEE and none for RR-PGEE. We stick to the more conservative threshold of 10 for the remainder of the analysis. The separated voxels seem to occur at the voxels with the lowest incidence rates, as might be expected (Figure B.3).

#### Interpretation

We inspect the total number of significant coefficients (FDR-corrected or thresholded at ±1.96) to quantify the spread of the effect of each covariate throughout white matter (see Table 6). The cross-sectional effect of age and CVR score have the most widely spread association with lesion probability with their longitudinal effects almost diminishing after FDR-correction. We further explore the effect of each covariate on lesion probability by plotting the raw *z*-scores obtained by the RR-PGEE marginal model (Figure 4); for the FDR-corrected results see Figure B.7 and Figures B.4 and B.5 for the equivalent RR-GEE results. The intra-subject correlation *α* and the dispersion parameter *ϕ* are also estimated voxel-wise (Figures 5 and B.6) revealing (i) positive within-subject correlation, and (ii) higher dispersion parameter estimates in areas of lower lesion incidence, i.e. since *ϕ* is a precision parameter, its higher values imply underdispersion at the boundaries.

**Table 6:**
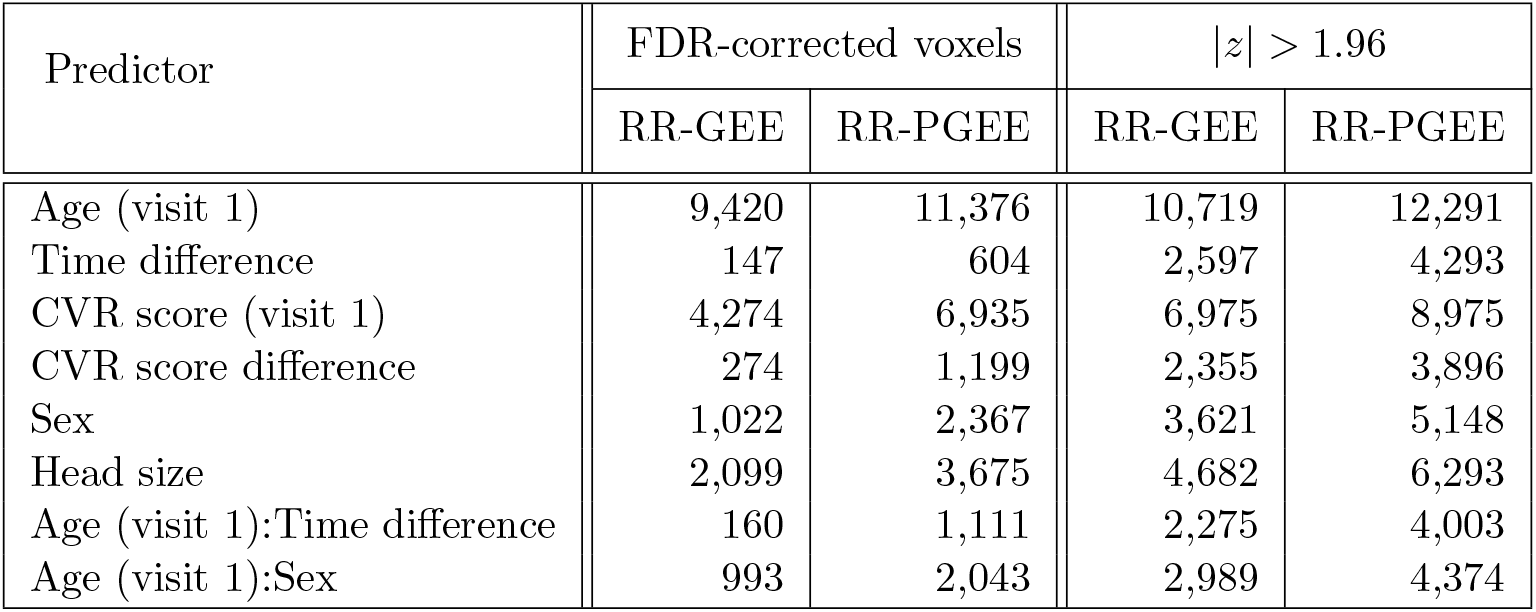
Number of significant voxels across predictors for RR-GEE and RR-PGEE estimates. Voxels with at least six individuals having a lesion across two visits explored (19,801 voxels in the brain mask). (Left) 5% FDR correction applied, i.e. number of FDR-corrected voxels is out of a total of 19,801 voxels, and (Right) fixed threshold of ±1.96 applied. Columns 2 and 3 complementary to Figures B.5 and B.7, respectively.

**Figure 4:**
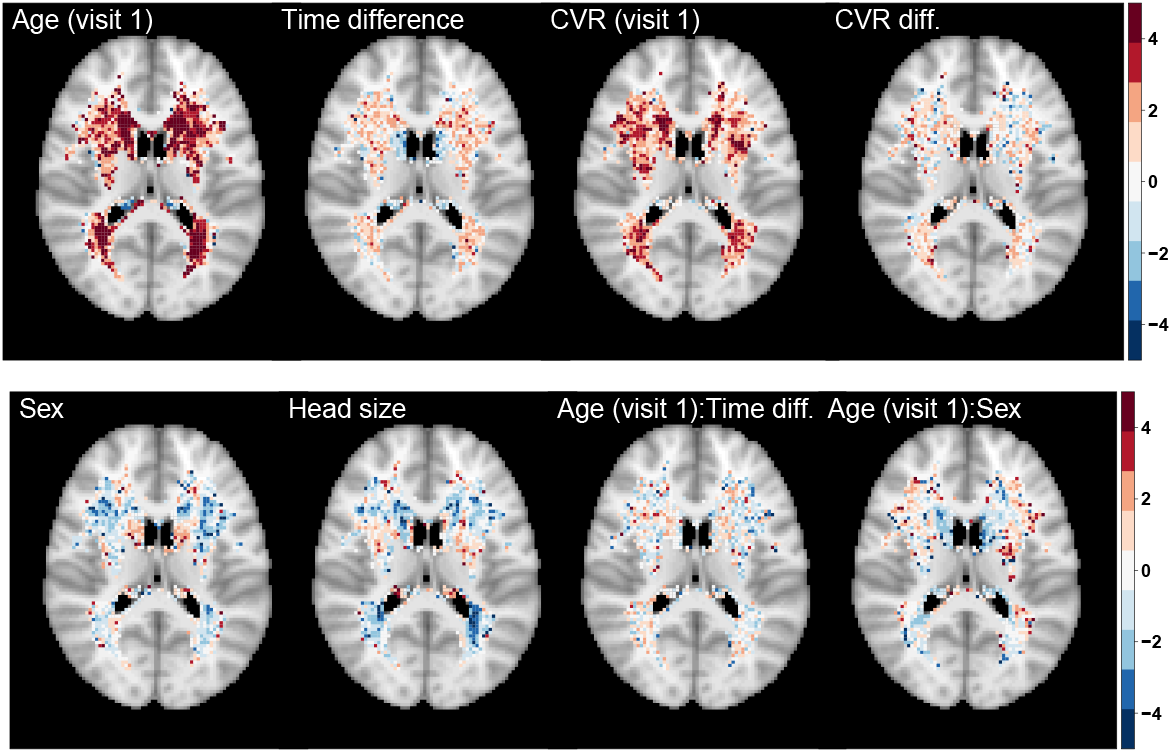
Significance maps (*z*-scores) based on RR-PGEE estimates 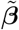. Data on 1,578 UKB participants across two visits and 19,801 voxels with six or more individuals having a lesion across two visits explored. Axial slice *z*=45 shown.

**Figure 5:**
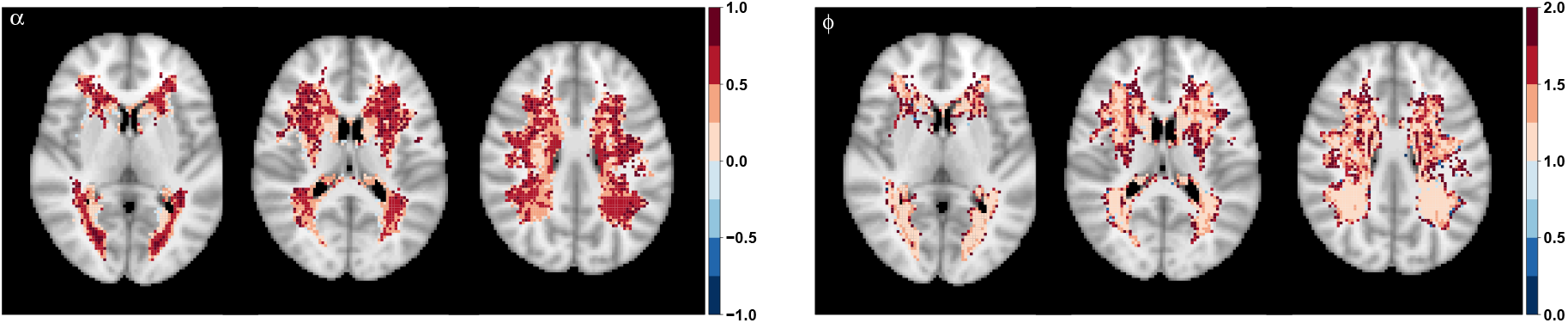
Correlation coefficient 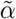 (left) and dispersion parameter 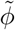 (right) based on fitting the marginal model using RR-PGEE. Data on 1,578 UKB participants across two visits and 19,801 voxels with six or more individuals having a lesion across two visits explored. Axial slices *z* = {40, 45, 50} shown.

Clearly the cross-sectional effects of age and CVR score are the most widely spread throughout white matter (Table 6, Figure 4) but *z*-scores suggest areas where we have detected strong associations between lesion probability and the covariates. The relative risks provide an insight into the magnitude of the effect. Without masking out any voxels, we plot the RR-PGEE relative risks for the four main covariates of interest in Figure 6. The baseline age relative risk suggests a remarkably uniform increase in lesion probability across the brain of about 10% (see Table 7 for relative risk summaries by empirical lesion incidence). In contrast, the baseline CVR relative risk suggests that an increase of 1 unit is associated with 36-39% increase in lesion probability in deep white matter where the lesion incidence is the lowest (up to 0.5%).

**Table 7:** Relative risks across empirical lesion incidence *p*_1_at visit 1 based on RR-PGEE estimates 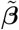. Mean and standard deviation (SD) are taken across voxels within each bin as defined in column 1. Data on 1,578 subjects across two visits and 19,801 regressions performed with the iterative algorithm failing for 2 voxels.

| $p_1$ | # voxels | Mean (SD) | | | | |
| --- | --- | --- | --- | --- | --- | --- |
| | | $p_2 - p_1$ | Relative risk $\exp(\tilde{\beta})$ | | | |
|  |  |  | Age<br>(visit 1) | Time<br>difference | CVR<br>(visit 1) | CVR<br>difference |
| [0; 0.0025) | 1,668 | 0.0017 (0.0010) | 1.11 (0.20) | 1.52 (2.02) | 1.39 (0.76) | 1.18 (0.75) |
| [0.0025; 0.005) | 6,232 | 0.0004 (0.0016) | 1.11 (1.04) | 1.07 (0.34) | 1.36 (0.49) | 1.19 (1.57) |
| [0.005; 0.01) | 4,810 | 0.0007 (0.0024) | 1.09 (0.09) | 1.05 (0.21) | 1.29 (0.28) | 1.08 (0.45) |
| [0.01; 1) | 7,089 | 0.0050 (0.0101) | 1.08 (0.06) | 1.05 (0.12) | 1.20 (0.16) | 1.07 (0.20) |
| [0; 1) | 19,799 | 0.0022 (0.0066) | 1.10 (0.59) | 1.09 (0.64) | 1.29 (0.40) | 1.12 (0.94) |

**Figure 6:**
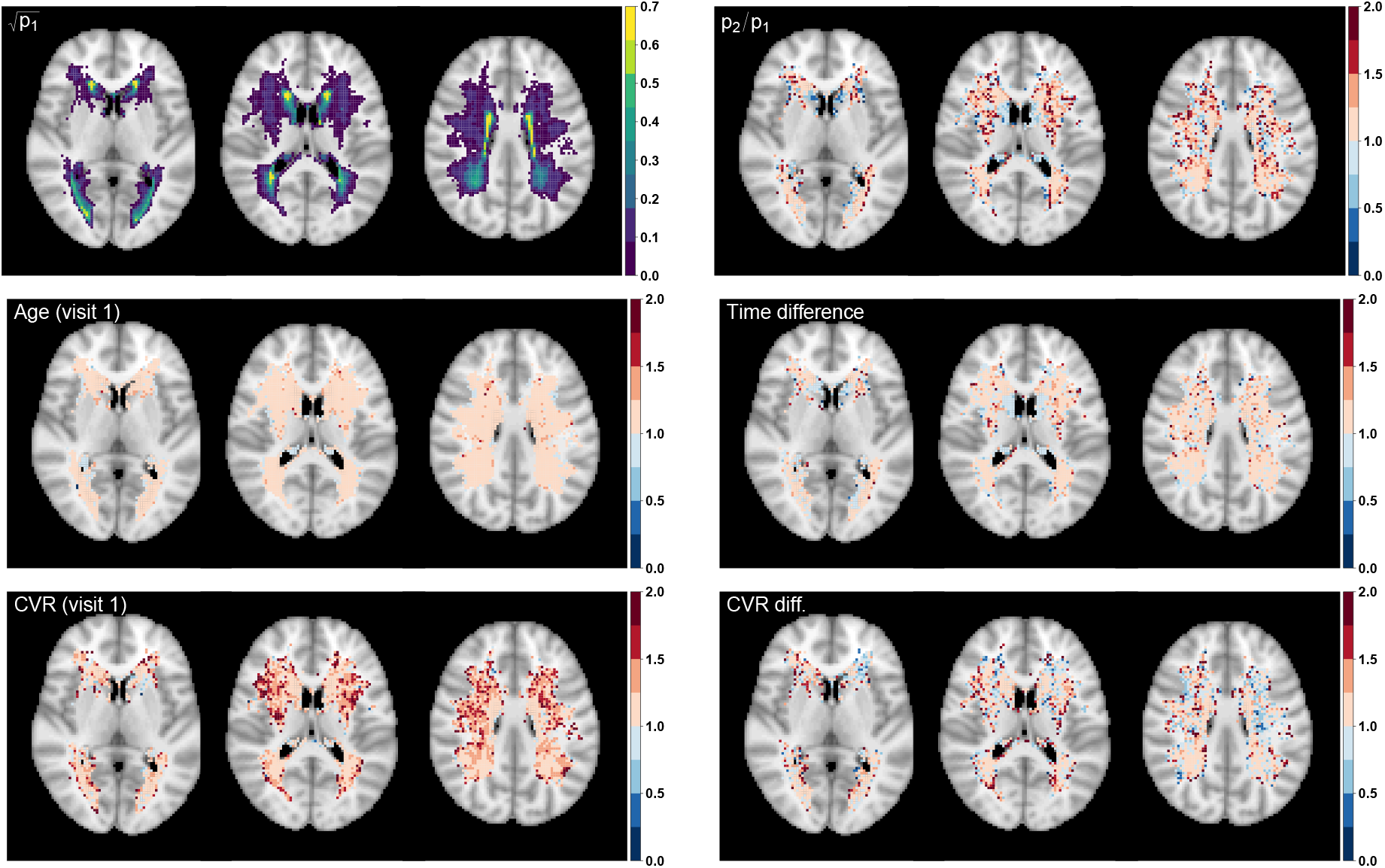
Relative risks exp 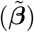 for four of the predictors in the RR-PGEE marginal model along with the square–root transformed empirical lesion incidence ***p***_1_ for visit 1 and the empirical relative risk ***p***_2_*/****p***_1_ (same as Figure 3) for reference. Data on 1,578 subjects across two visits and 19,801 voxels explored. Axial slices *z* = {40, 45, 50} shown.

#### Relative risk vs odds ratio modelling

We further explore the relative risks obtained from relative risk regression (RR-GEE and RR-PGEE) and from logit-link GEE (OR-PGEE method by Mondol and Rahman 2019). Figure 7 explores the obtained relative risks for age at visit 1 and it reveals that (i) both penalized GEE methods result in shrinkage of the relative risks towards 1 when compared to the RR-GEE, i.e. shrinkage of the estimated coefficients towards 0, (ii) applying a penalty to the log-odds or to the log-relative risks seems to lead to largely similar results.

**Figure 7:**
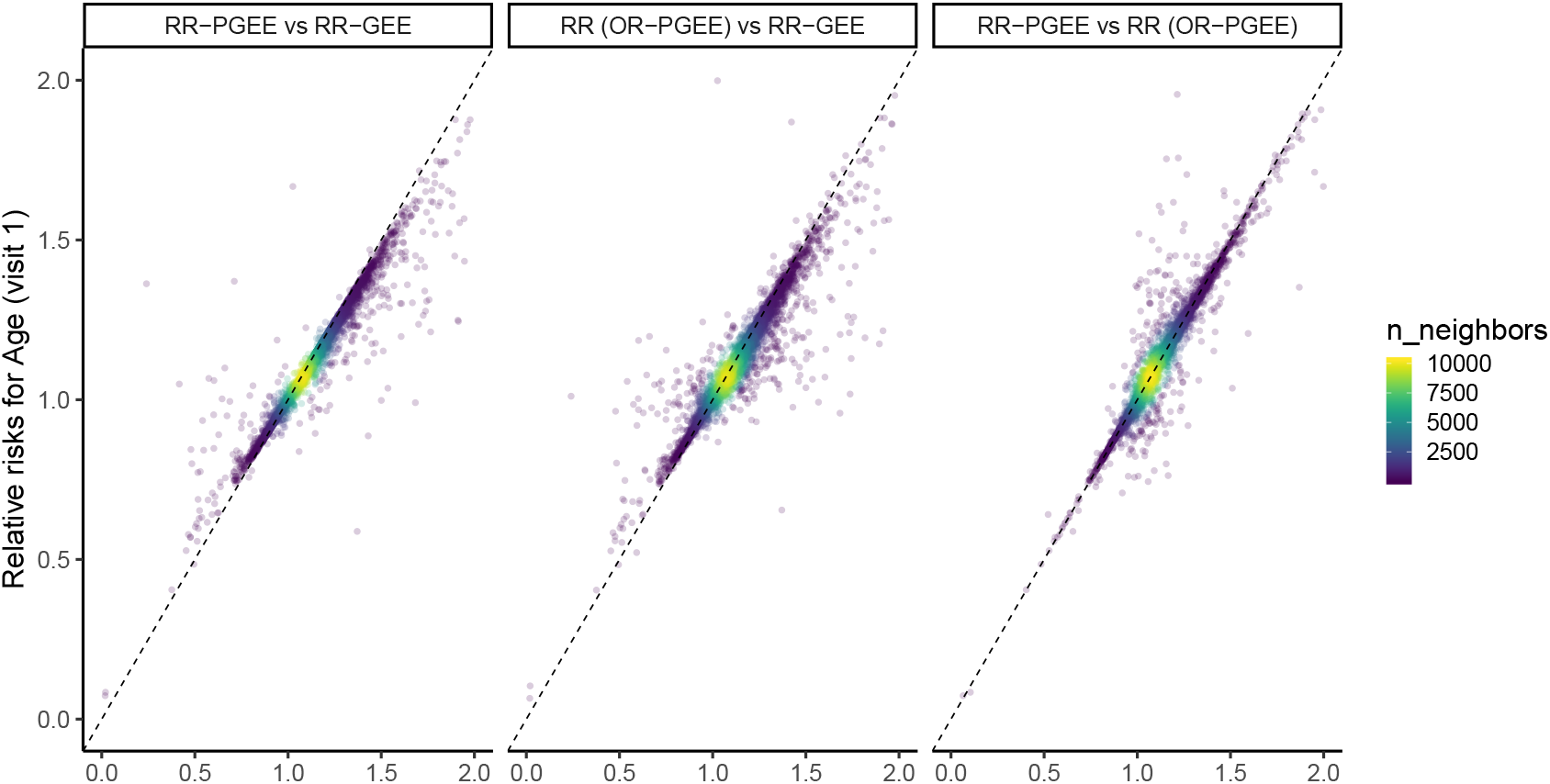
Relative risk for age at visit 1 obtained across the three methods. The title of each panel is in the form ‘y vs x’, i.e. relative risks for age obtained by fitting RR-PGEE plotted against relative risks obtained by fitting RR-GEE. RR (OR-PGEE) are the transformed relative risks resulting from the odds ratios (OR (OR-PGEE)) obtained by fitting OR-PGEE. Data on 1,578 subjects across two visits and 19,801 voxels explored.

#### Performance

We can obtain in-sample predictions voxel-wise 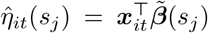 across subjects *i* and and visits *t* using the RR-GEE estimates 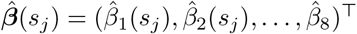 for the intercept and the eight explanatory variables as in (8). Similarly, we obtain voxel-wise predictions 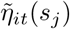 using RR-PGEE estimates 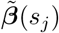 across subjects *i* and visits *t*. Note, we have *N* × *T* = 1578 × 2 = 3, 156 predicted maps for each method with predicted linear predictor values spread across 19,801 voxels.

We then find that all 3,156 maps have at least one non-negative value 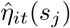 for at least one voxel *s*_*j*_ for given subject *i* and time point *t*) using the RR-GEE estimates (if we do not exclude voxels with boundary estimates) and 1,731 maps (55%) using RR-PGEE estimates, respectively. However, in the majority of predicted maps, where we observe non-negative prediction values, only up to ten voxels out of about 20,000 voxels run into this issue (71% out of 3,156 maps using RR-GEE estimates and 73% out of 1,733 maps using RR-PGEE estimates). Note that the penalty we introduce in the estimating equations does not ensure the predicted values result in valid probability values, but its use seems to alleviate the problem.

#### Implementation details

Performing 19,801 regressions for all three methods - RR-GEE, RR-PGEE and OR-PGEE - with data on 1,578 subjects at two time points would take long to perform serially. Our approach is to splits the total number of regressions into subsets of 500 and then execute each subset of 500 regressions as a single-core job, i.e. a total of 40 jobs running in parallel for each of the methods. Our implementation of RR-GEE and RR-PGEE is entirely in `R`as is the OR-PGEE one by Mondol and Rahman (2019). Each job of 500 regressions takes about 50-60 minutes for RR-GEE, 2-2.5 hours for RR-PGEE and 3-4 hours for OR-PGEE. Note that our code is not optimised and, in particular, includes computation of boundary estimates criterion values at each iteration that would not be needed in routine usage.

## 5. Discussion

Taking a marginal approach to modelling correlated binary data, in this paper we have introduced penalized generalized estimating equations for relative risk regression (RR-PGEE). Our work was motivated by binary brain lesion data derived from MRI scans, since brain lesions have varying incidence across the brain. As a result, odds ratios estimated from GEE with logistic regression structures cannot always safely approximate risk ratios. On the other hand, use of log-link regression structures with the binomial variance function may lead to estimation instabilities when event probabilities are close to 1. To obtain finite estimates when dealing with rare outcomes or small sample size, we introduced a Jeffreys-prior penalty to the GEE for relative risk regression in a similar manner to bias-reduction methods in GLMs, while using the identity variance function and unknown dispersion for extra stability of the estimates.

We tested the performance of the RR-PGEE estimates through extensive simulations. The penalized approach provided finite estimates and achieved convergence even for simulated data sets where RR-GEE showed signs of separation though estimates diverging to infinity and lack of convergence, or the iterative estimation procedure failed. The inclusion of the penalty also offered some reduction in the mean squared error of the estimates when boundary estimates were not present, with the bias being comparable between the two approaches.

Applying the alternative modelling approaches to a subset of the UK Biobank data, we explored the association between brain lesions, and ageing and cerebrovascular risk. The RR-PGEE approach resulted in stable estimates across the entire brain with RR-GEE resulting in about 5% missing or diverging estimates. The alternative penalized logit-link GEE (Mondol and Rahman, 2019) resulted in highly similar estimates (transforming the odds ratios to risk ratios).

The longitudinal effects of age and CVR score did not result in largely significant estimates across the brain, but this replicates what has been previously shown in the clinical literature (Sachdev et al., 2007). CVR factors and age are undoubtedly strong predictors of lesion incidence and total burden of lesions when measured cross-sectionally. In contrast, baseline total lesion load is a better predictor of lesion incidence, than CVR burden or age, when measured longitudinally. The cross-sectional effect of CVR score showed an interesting localized effect in deep white matter, which is likely the result of small vessel disease. The increase in deep white matter lesion probability associated with CVR reflects the impact of CVR factors like hypertension on small vessels. Also, age seemed to have almost homogeneous effect on lesion probability across the brain suggesting increased risk of lesion occurrence with ageing.

### Limitations

Throughout the simulation scenarios, we observed that the sandwich variance slightly underestimated the true variance when using RR-PGEE. In small samples, the sandwich variance is known to underestimate the true variance and thus small-sample bias-correction to the sandwich estimator is considered in the literature (Fay and Graubard, 2001; Mancl and DeRouen, 2001; Morel et al., 2003). Studying appropriate small-sample sandwich estimator adjustments is out of the scope of this work and we believe it does not impact the big data application that motivated our own work. Non-convergence could be due to the choice of the starting values for the IRLS algorithm and more elaborate testing schemes for starting values can result in convergence for the data sets where the boundary estimates criterion was lower than the selected threshold but both methods did not converge.

When exploring the relative risks arising from the UKB data analysis, we do not restrict our attention to voxels passing FDR-correction. We emphasize on the smoothness of the relative risk maps and on the suggested patterns, but we mostly want to highlight the potential of the method and the usefulness of those maps. Surely, clinical interpretations should be approached with extra caution.

We observe that the empirical lesion risk around the ventricles is lower than 1, suggesting that some of the lesions disappear over time. We believe that this phenomenon is not biological but could rather be explained by a mixture of the following: the lesion segmentation is leading to more false positives when lesion load is low (younger subjects), the spatial normalisation to MNI is not optimized for white matter, the caudate is known to be shrinking with age and ventricles are known to enlarge with age which might lead to some registration challenges too. None of those potential segmentation and registration challenges should have affected the modelling approach we suggested, we just note that the estimates in the periventricular areas should be interpreted with caution.

One of the main disadvantages of RR-GEE and RR-PGEE is that they do not guarantee the predicted probabilities would lie in the 0-1 range. If the linear predictor is greater than 0, the resulting predicted probability would be above 1. This did happen very rarely for the simulated data sets but it happened more often for the noisier real data set for single voxels. The main aim of the current work was to provide stable and interpretable estimates, i.e. inference, as opposed to prediction. We have demonstrated that the estimated relative risks were largely similar to the relative risks obtained from logistic regression (transforming odds ratios to risk ratios) ensuring that inference was not affected by the unsupported predicted values.

## Acknowledgments

We are grateful to Ludovica Griffanti and Stephen Smith for their valuable input on the lesion segmentation and registration practices in the UK Biobank. The computational aspects of this research were supported by the Wellcome Trust Core Award Grant Number 203141/Z/16/Z and the NIHR Oxford BRC. The views expressed are those of the author(s) and not necessarily those of the NHS, the NIHR or the Department of Health.

## Data availability statement

The `R` code used to fit the marginal models using RR-GEE and RR-PGEE is available at the GitHub repository https://github.com/petyakindalova/PGEE_Illustration along with an illustration of the code used for the simulation study. The spatial relative risk maps produced as part of the real data application are available at NeuroVault https://neurovault.org/collections/SUDHNHAA/.

## A. Derivation of penalty term

To obtain the form of the Jeffreys-prior penalty term *A*_*p*_(***β, α***) for this RR-GEE (Equation (2)), we first take the derivative of *U* wrt *β*_*p*_ by using the product and chain rules:

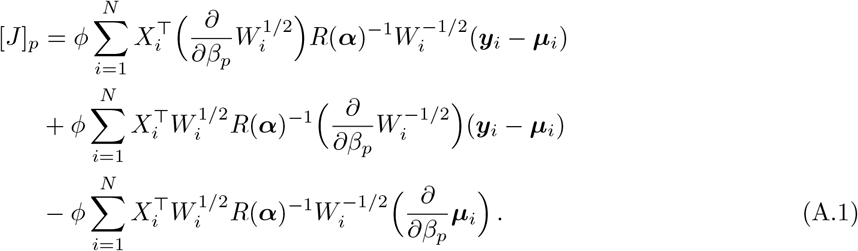

Since E(***y***_*i*_) = ***µ***_*i*_ by assumption and ***y***_*i*_ − ***µ***_*i*_ come linearly into (A.1), the *p*-th row of the expected negative Hessian matrix *I* is

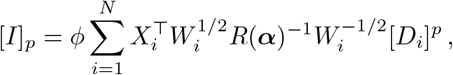

where [*D*_*i*_]^*p*^ is the *p*-th column of *D*_*i*_. So,

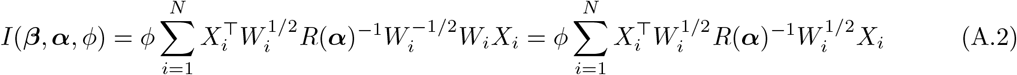

We further take the derivative of *I* and by using the product and chain rules we get

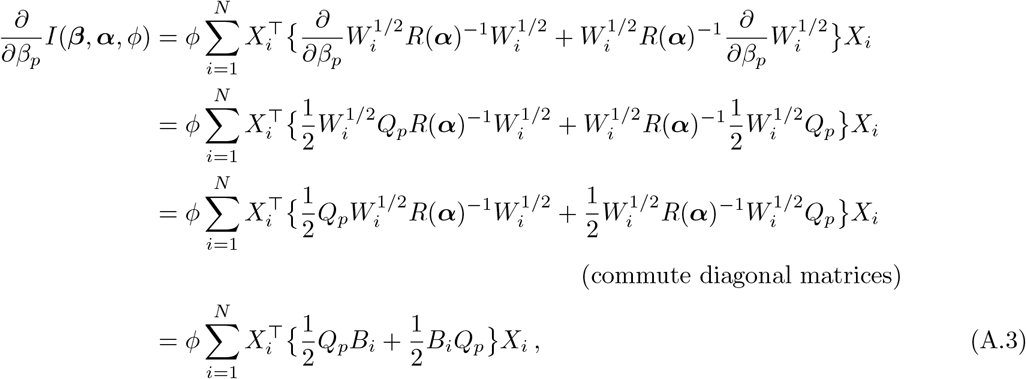

where *Q*_*p*_ = diag(*x*_*i*1*p*_, …, *x*_*iT p*_) and 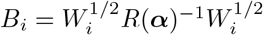. Combining expressions (A.2) and (A.3) we construct the penalty term:

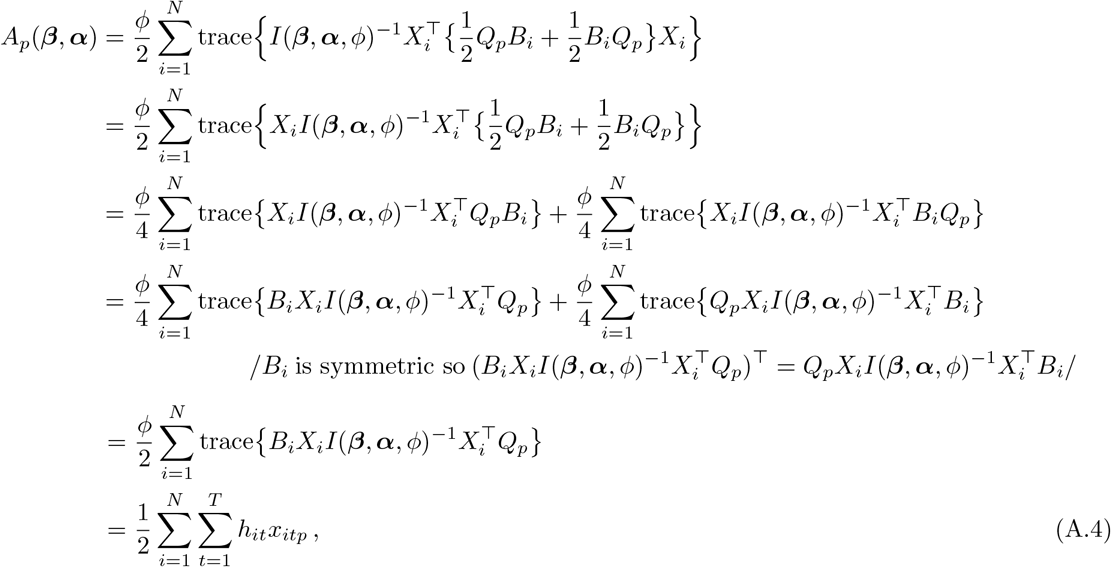

where *h*_*it*_ is the *t*-th diagonal element of the *i*-th block of the projection matrix

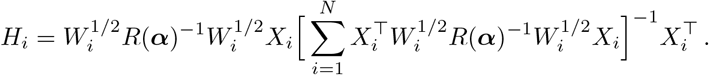

## B. Complementary tables and figures

**Figure B.1:**
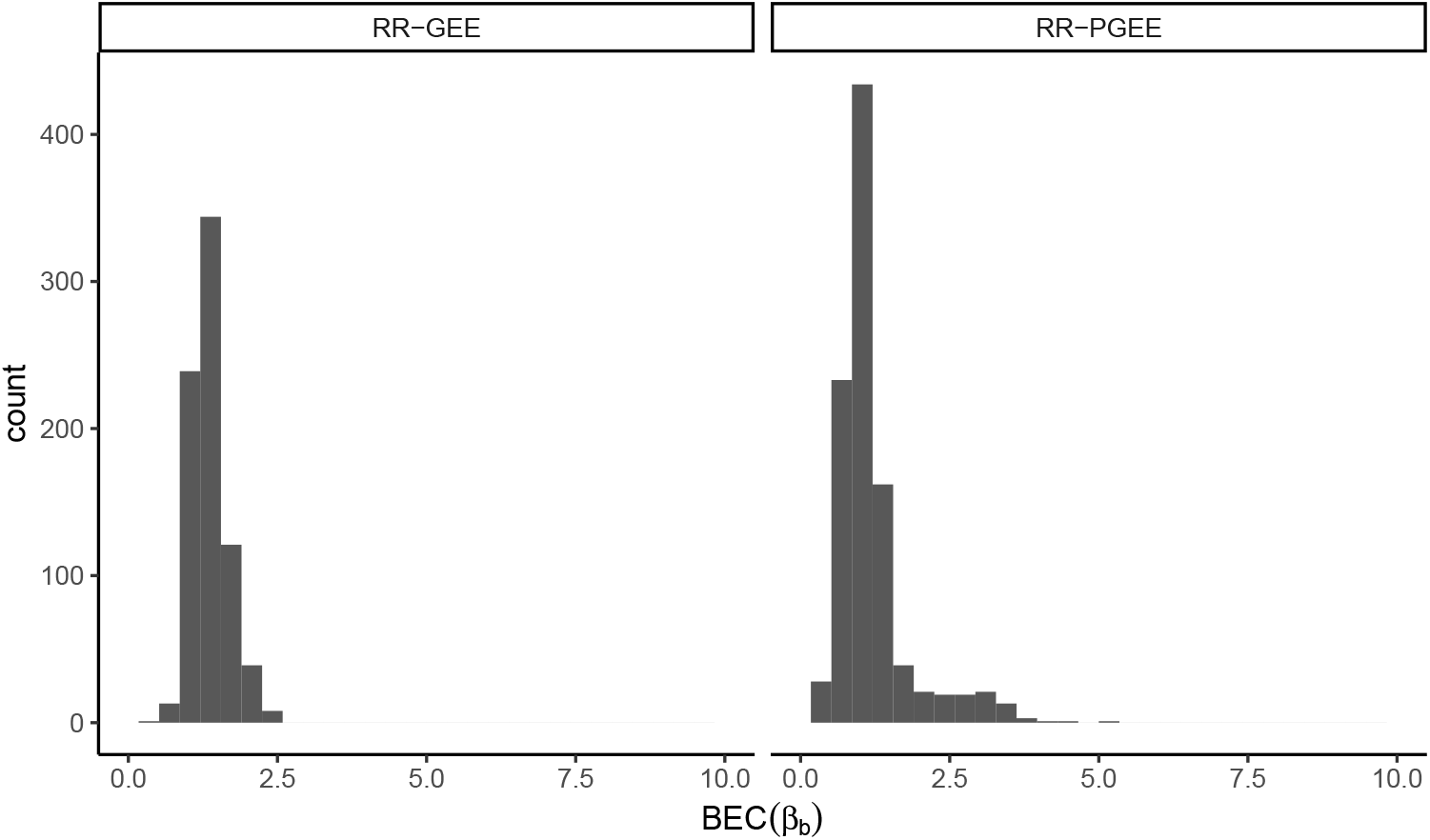
Boundary estimates criterion threshold is empirically selected and set to 10 for the simulation study. (Left) (BÊC_2_) and (Right) 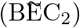 across 1,000 simulated data sets from the base simulation scenario; parameters are set to ***β*** = (*β*_1_, *β*_*b*_, *β*_*c*_)^⊤^ = (−4, 1.6, 0.2)^⊤^, *N* = 50, *c* = 0.2, *α* = 0.4 and all existing BEC values are shown for the fixed *x*-axis limits.

**Table B.1:** Bias and mean-squared error of the binary covariate coefficient *β*_*b*_ across simulation scenarios. The simulation parameters are set to ***β*** = (*β*_1_, *β*_*b*_, *β*_*c*_)^⊤^ = (−4, 1.6, 0.2)^⊤^, *α* = 0.4 and each row of the table summarizes 1,000 replications with just one parameter altered, e.g. the first block of rows represent scenarios when *c* is altered for sample size *N* = 100 and for sample size *N* = 500, respectively. Summaries included are conditional on both estimation methods converging and the estimates being finite (no boundary estimates), with unconditional summaries for RR-PGEE included in parenthesis.

| $N = 100$ | | Finite & converging | | | | |
| --- | --- | --- | --- | --- | --- | --- |
| Parameter | Value | N.sim | $B(\hat{\beta}_b)$ | $B(\tilde{\beta}_b)$ | $MSE(\hat{\beta}_b)$ | $MSE(\tilde{\beta}_b)$ |
| $c$ | **0.2** | 963 (995) | 0.05 | 0.03 (0.07) | 0.75 | 0.56 (0.82) |
|  | 0.3 | 957 (1000) | 0.11 | 0.02 (0.12) | 0.66 | 0.49 (0.86) |
|  | 0.4 | 941 (1000) | 0.10 | -0.04 (0.11) | 0.67 | 0.50 (0.85) |
|  | 0.5 | 906 (1000) | 0.07 | -0.11 (0.10) | 0.60 | 0.44 (0.84) |
|  | 0.6 | 840 (1000) | 0.03 | -0.19 (0.11) | 0.57 | 0.44 (0.85) |
|  | 0.7 | 760 (1000) | -0.10 | -0.37 (0.02) | 0.53 | 0.49 (0.79) |
|  | 0.8 | 615 (1000) | -0.29 | -0.63 (-0.12) | 0.55 | 0.69 (0.66) |
| $N = 500$ | | Finite & converging | | | | |
| Parameter | Value | N.sim | $B(\hat{\beta}_b)$ | $B(\tilde{\beta}_b)$ | $MSE(\hat{\beta}_b)$ | $MSE(\tilde{\beta}_b)$ |
| $c$ | **0.2** | 1000 (1000) | $7 \times 10^{-4}$ | -0.01 (-0.01) | 0.13 | 0.12 (0.12) |
|  | 0.3 | 1000 (1000) | 0.01 | -0.01 (-0.01) | 0.11 | 0.11 (0.11) |
|  | 0.4 | 1000 (1000) | 0.02 | -0.01 (-0.01) | 0.11 | 0.11 (0.11) |
|  | 0.5 | 1000 (1000) | 0.03 | -0.02 (-0.02) | 0.15 | 0.13 (0.13) |
|  | 0.6 | 1000 (1000) | 0.08 | 0.02 (0.02) | 0.19 | 0.16 (0.16) |
|  | 0.7 | 999 (1000) | 0.11 | 0.03 (0.03) | 0.28 | 0.22 (0.23) |
|  | 0.8 | 993 (1000) | 0.18 | 0.03 (0.04) | 0.45 | 0.30 (0.34) |

B: bias; MSE: mean squared error.

**Figure B.2:**
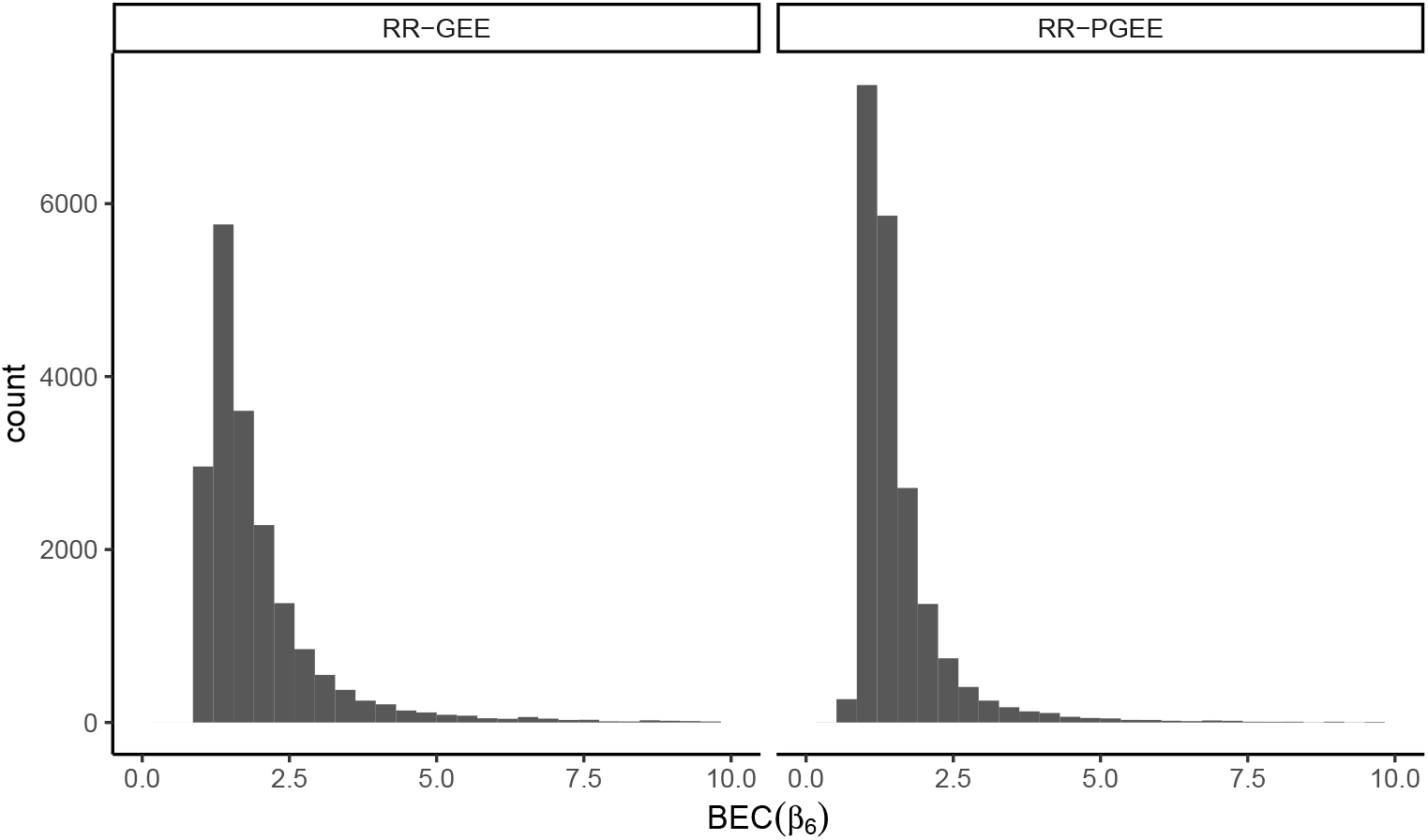
Boundary estimates criterion threshold is empirically selected and set to 10 for UKB analysis and the BEC values are shown for the sex covariate. (Left) (BÊC_6_) and (Right) 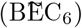 across 19,801 voxels; all existing BEC values are shown for the fixed *x*-axis limits. Voxels not shown on the histograms include 763 voxels with BEC values greater than 10 for any of the 9 covariates for RR-GEE and 90 for RR-PGEE, respectively, as well as any missing values due to IRLS failure.

**Table B.2:** List of codes of the UK Biobank Data-field ‘Non-cancel illness’ (http://biobank.ndph.ox.ac.uk/showcase/field.cgi?id=20002) used for exclusion of participants in the data cleaning process. 118 illnesses reported across 106 participants.

| Coding | Description | Number of participants |
| --- | --- | --- |
| 1081 | Stroke | 29 |
| 1082 | Transient ischaemic attack | 18 |
| 1083 | Subdural haematoma/haemorrhage | 2 |
| 1086 | Subarachnoid haemorrhage | 0 |
| 1240 | Neurological injury/trauma | 2 |
| 1243 | Psychological/psychiatric problem | 0 |
| 1244 | Infection of the nervous system | 0 |
| 1245 | Brain abscess/Intracranial abscess | 1 |
| 1246 | Encephalitis | 1 |
| 1247 | Meningitis | 14 |
| 1258 | Chronic neurological problem | 0 |
| 1259 | Motor Neuron Disease | 1 |
| 1261 | Multiple Sclerosis | 10 |
| 1262 | Parkinson’s disease | 4 |
| 1263 | Dementia/Alzheimer’s disease/Cognitive impairment | 1 |
| 1264 | Epilepsy | 13 |
| 1266 | Head Injury | 4 |
| 1408 | Alcoholism | 1 |
| 1409 | Opioid dependency | 0 |
| 1410 | Other dependency | 0 |
| 1434 | Other neurological problem | 6 |
| 1491 | Brain haemorrhage | 5 |
| 1583 | Ischaemic stroke | 1 |
| 1626 | Fracture skull/head | 5 |

**Figure B.3:**
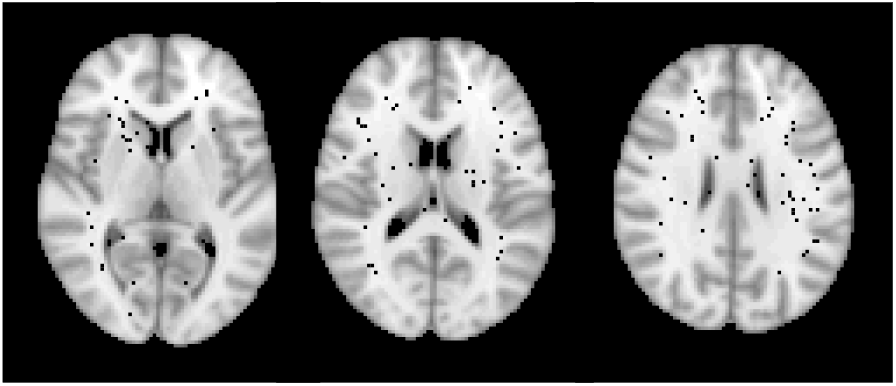
Voxels with boundary estimates in black, where about 5% of voxels have boundary estimates criterion values higher than 10 or missing values due to failure of the IRLS algorithm. Voxels with boundary estimates seem to occur in regions of low lesion incidence. Axial slices *z* = {40, 45, 50} shown.

**Figure B.4:**
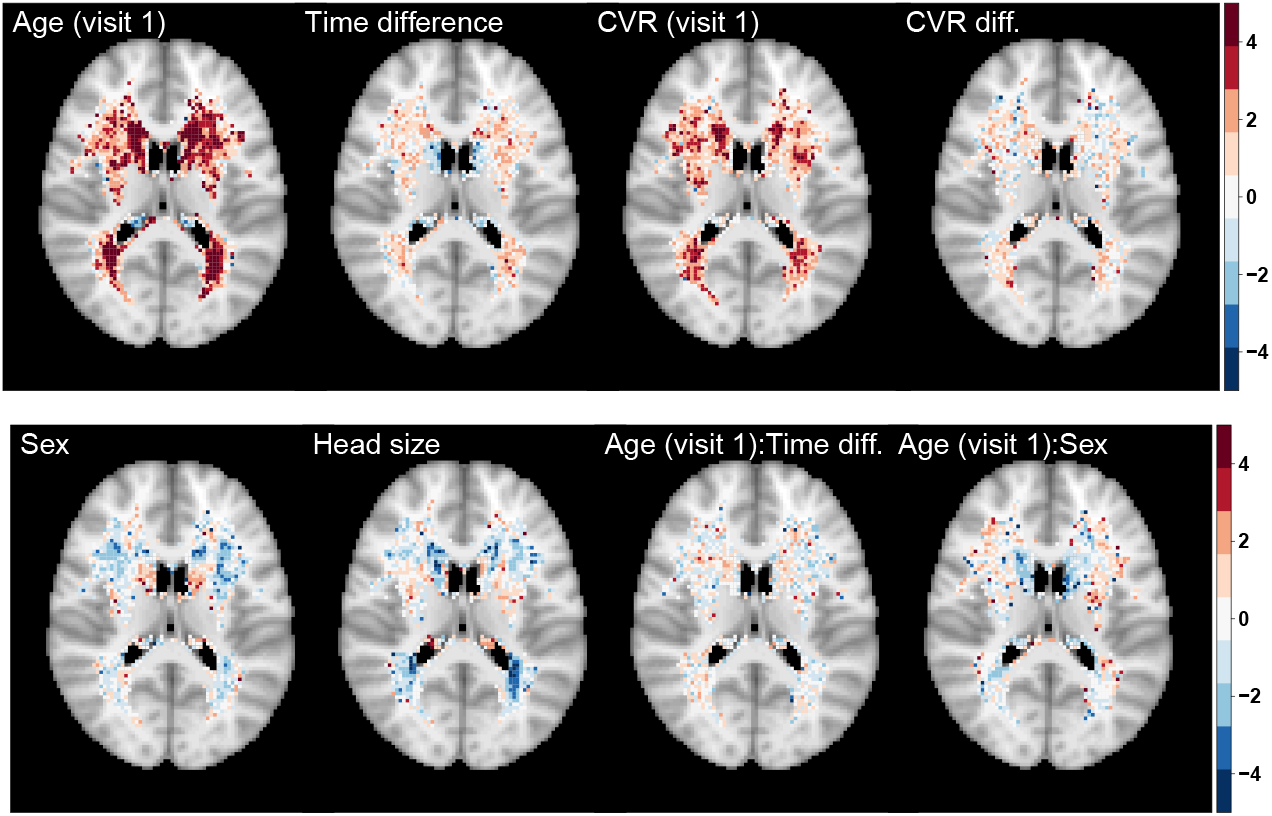
Significance maps (*z*-scores) based on RR-GEE estimates 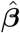. Data on 1,578 UKB participants across two visits and 19,801 voxels with six or more individuals having a lesion across two visits explored. Axial slice *z* = 45 shown.

**Figure B.5:**
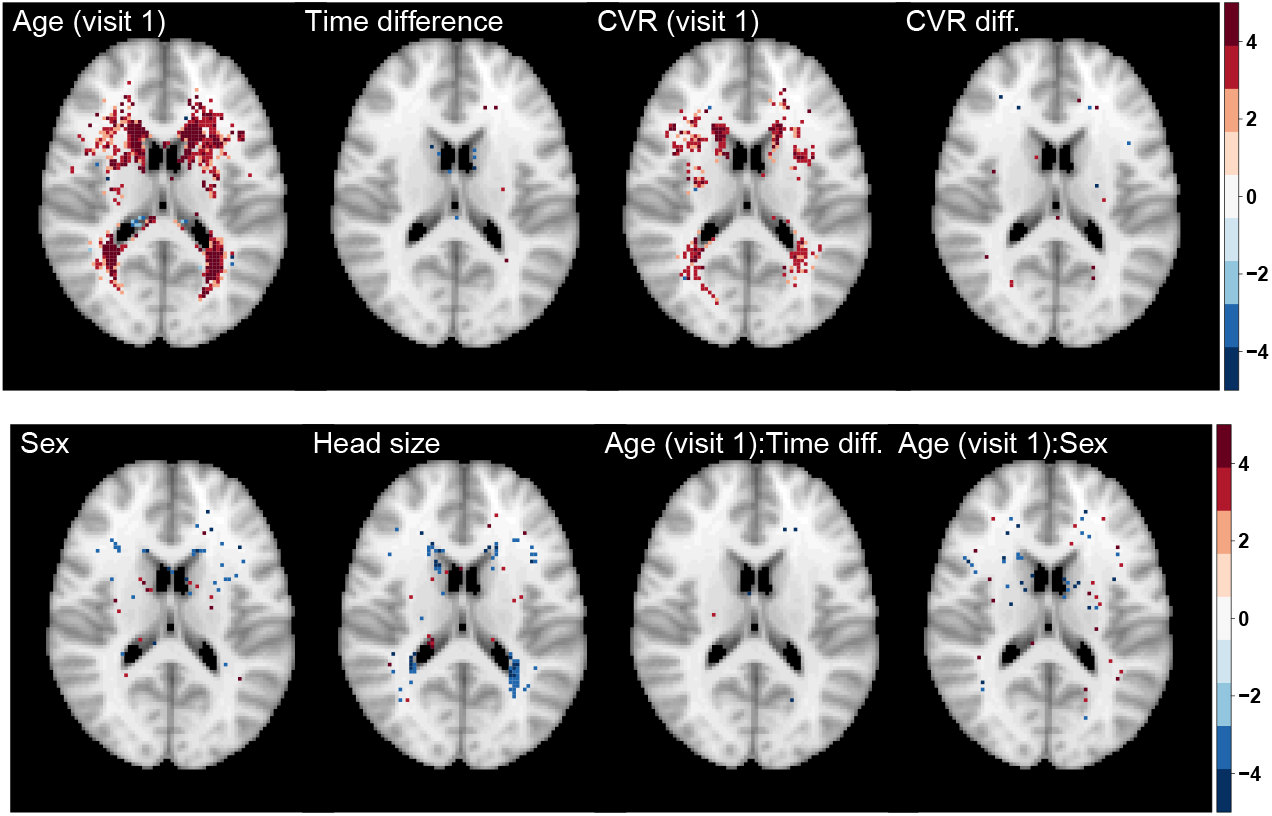
Significance maps (*z*-scores) based on RR-GEE estimates 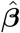. Data on 1,578 UKB participants across two visits and 19,801 voxels with six or more individuals having a lesion across two visits explored; 5%-FDR correction applied. Axial slice *z* = 45 shown.

**Figure B.6:**
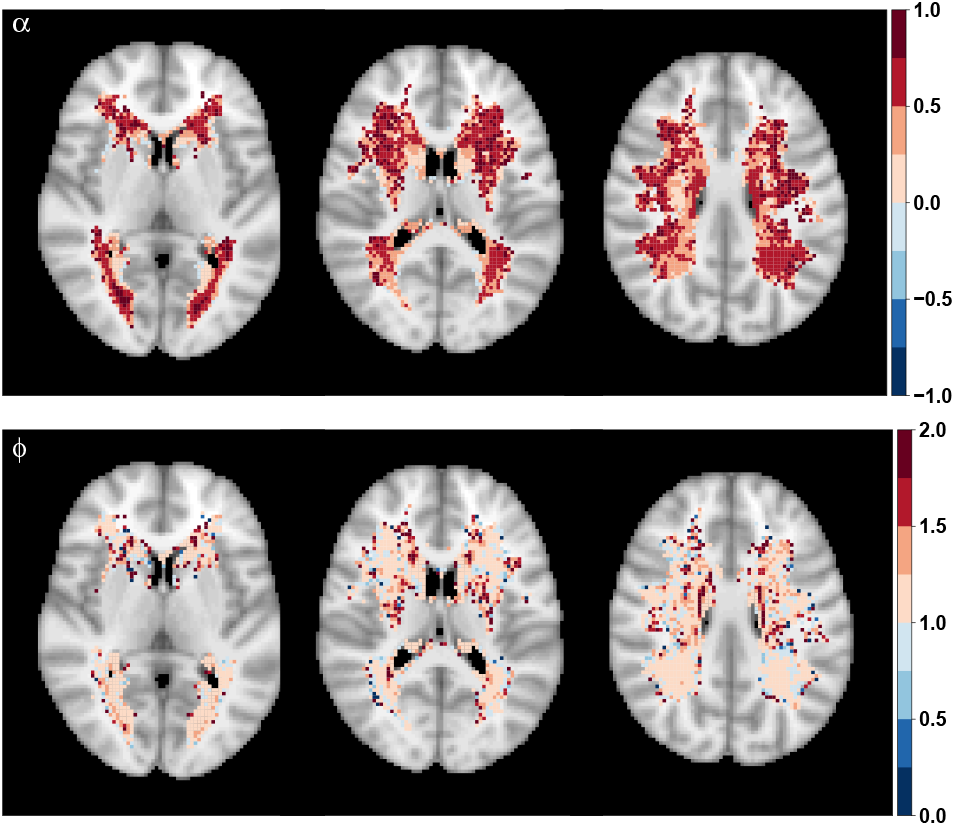
Correlation coefficient 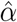 (top) and dispersion parameter 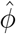 (bottom) based on fitting the marginal model using RR-GEE. Data on 1,578 UKB participants across two visits and 19,801 voxels with six or more individuals having a lesion across two visits explored. Axial slices *z* = {40, 45, 50} shown.

**Figure B.7:**
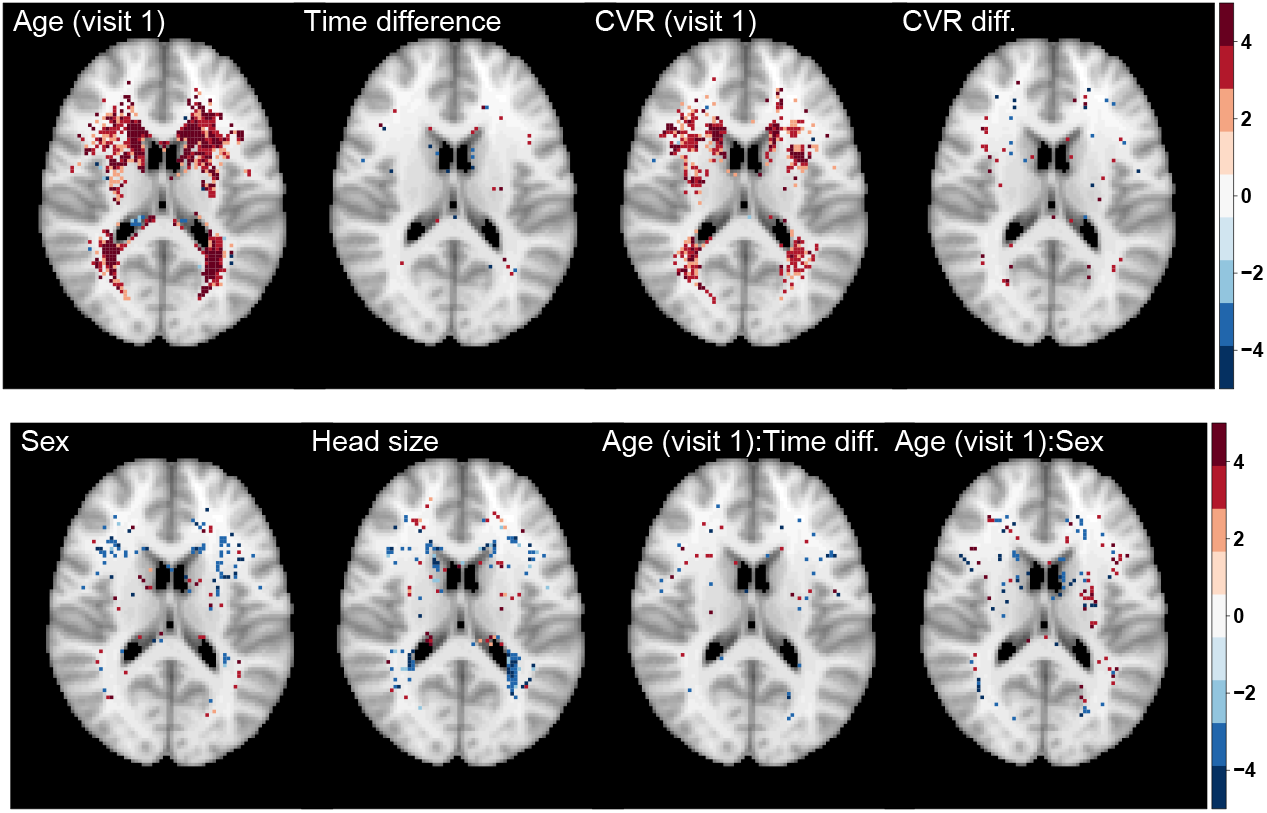
Significance maps (*z*-scores) based on RR-PGEE estimates 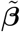. Data on 1,578 UKB participants across two visits and 19,801 voxels with six or more individuals having a lesion across two visits explored; 5%-FDR correction applied. Axial slice *z* = 45 shown.

## Footnotes

http://biobank.ndph.ox.ac.uk/showcase/search.cgi

## References

Albert, A. and Anderson, J. A. (1984). On the existence of maximum likelihood estimates in logistic regression models. Biometrika, 71(1).

Alfaro-Almagro, F., Jenkinson, M., Bangerter, N. K., Andersson, J. L., Griffanti, L., Douaud, G., Sotiropoulos, S. N., Jbabdi, S., Hernandez-Fernandez, M., Vallee, E., Vidaurre, D., Webster, M., McCarthy, P., Rorden, C., Daducci, A., Alexander, D. C., Zhang, H., Dragonu, I., Matthews, P. M., Miller, K. L., and Smith, S. M. (2018). Image processing and Quality Control for the first 10,000 brain imaging datasets from UK Biobank. NeuroImage, 166.

Alfaro-Almagro, F., McCarthy, P., Afyouni, S., Andersson, J. L., Bastiani, M., Miller, K. L., Nichols, T. E., and Smith, S. M. (2020). Confound modelling in UK Biobank brain imaging. NeuroImage, 224:117002.

Benjamini, Y. and Hochberg, Y. (1995). Controlling the False Discovery Rate: A Practical and Powerful Approach to Multiple Testing. Journal of the Royal Statistical Society: Series B (Methodological), 57(1).

Carter, R. E., Lipsitz, S. R., and Tilley, B. C. (2005). Quasi-likelihood estimation for relative risk regression models. Biostatistics, 6(1).

Fay, M. P. and Graubard, B. I. (2001). Small-sample adjustments for Wald-type tests using sandwich estimators. Biometrics, 57(4).

Firth, D. (1993). Bias Reduction of Maximum Likelihood Estimates. Biometrika, 80(1).

Fitzmaurice, G., Laird, N., and Ware, J. (2011). Applied longitudinal Analysis (2nd Edition). John Wiley & Sons, Hoboken.

Fitzmaurice, G. M. (1995). A Caveat Concerning Independence Estimating Equations with Multivariate Binary Data. Biometrics, 51(1).

Fitzmaurice, G. M., Lipsitz, S. R., Arriaga, A., Sinha, D., Greenberg, C., and Gawande, A. A. (2014). Almost efficient estimation of relative risk regression. Biostatistics, 15(4).

Gardiner, J. C., Luo, Z., and Roman, L. A. (2009). Fixed effects, random effects and GEE: What are the differences? Statistics in Medicine, 28(2).

Greenland, S. and Thomas, D. C. (1982). On the need for the rare disease assumption in case-control studies. American Journal of Epidemiology, 116(3).

Greenland, S., Thomas, D. C., and Morgenstern, H. (1986). The rare-disease assumption revisited: A critique of ”estimators of relative risk for case-control studies”. American Journal of Epidemiology, 124(6).

Griffanti, L., Zamboni, G., Khan, A., Li, L., Bonifacio, G., Sundaresan, V., Schulz, U. G., Kuker, W., Battaglini, M., Rothwell, P. M., and Jenkinson, M. (2016). BIANCA (Brain Intensity AbNormality Classification Algorithm): A new tool for automated segmentation of white matter hyperintensities. NeuroImage, 141.

Heagerty, P. J. and Zeger, S. L. (2000). Marginalized multilevel models and likelihood inference. Statistical Science, 15(1).

Hubbard, A. E., Ahern, J., Fleischer, N. L., Laan, M. V. D., Lippman, S. A., Jewell, N., Bruckner, T., and Satariano, W. A. (2010). To GEE or not to GEE: Comparing population average and mixed models for estimating the associations between neighborhood risk factors and health. Epidemiology, 21(4).

Knol, M. J., Le Cessie, S., Algra, A., Vandenbroucke, J. P., and Groenwold, R. H. (2012). Overestimation of risk ratios by odds ratios in trials and cohort studies: Alternatives to logistic regression. CMAJ, 184(8).

Knol, M. J., Vandenbroucke, J. P., Scott, P., and Egger, M. (2008). What do case-control studies estimate? Survey of methods and assumptions in published case-control research. American Journal of Epidemiology, 168(9).

Kosmidis, I. and Firth, D. (2021). Jeffreys-prior penalty, finiteness and shrinkage in binomial-response generalized linear models. Biometrika, 108(1).

Laird, N. M. and Ware, J. H. (1982). Random-Effects Models for Longitudinal Data. Biometrics, 38(4).

Lampe, L., Kharabian-Masouleh, S., Kynast, J., Arelin, K., Steele, C. J., Löffler, M., Witte, A. V., Schroeter, M. L., Villringer, A., and Bazin, P. L. (2019). Lesion location matters: The relationships between white matter hyperintensities on cognition in the healthy elderly. Journal of Cerebral Blood Flow and Metabolism, 39(1):36–43.

Lee, Y. and Nelder, J. A. (2004). Conditional and marginal models: Another view. Statistical Science, 19(2).

Liang, K. Y. and Zeger, S. L. (1986). Longitudinal data analysis using generalized linear models. Biometrika, 73(1).

Lindsey, J. K. and Lambert, P. (1998). On the appropriateness of marginal models for repeated measurements in clinical trials. Statistics in Medicine, 17(4).

Litière, S., Alonso, A., and Molenberghs, G. (2007). Type I and type II error under random-effects misspecification in generalized linear mixed models. Biometrics, 63(4).

Mancl, L. A. and DeRouen, T. A. (2001). A covariance estimator for GEE with improved small-sample properties. Biometrics, 57(1).

Mansournia, M. A., Geroldinger, A., Greenland, S., and Heinze, G. (2018). Separation in Logistic Regression: Causes, Consequences, and Control. American Journal of Epidemiology, 187(4).

McNutt, L. A., Wu, C., Xue, X., and Hafner, J. P. (2003). Estimating the relative risk in cohort studies and clinical trials of common outcomes. American Journal of Epidemiology, 157(10).

Miller, K. L., Alfaro-Almagro, F., Bangerter, N. K., Thomas, D. L., Yacoub, E., Xu, J., Bartsch, A. J., Jbabdi, S., Sotiropoulos, S. N., Andersson, J. L., Griffanti, L., Douaud, G., Okell, T. W., Weale, P., Dragonu, I., Garratt, S., Hudson, S., Collins, R., Jenkinson, M., Matthews, P. M., and Smith, S. M. (2016). Multimodal population brain imaging in the UK Biobank prospective epidemiological study. Nature Neuroscience, 19(11).

Mondol, M. H. and Rahman, M. S. (2019). Bias-reduced and separation-proof GEE with small or sparse longitudinal binary data. Statistics in Medicine, 38(14).

Morel, J. G., Bokossa, M. C., and Neerchal, N. K. (2003). Small sample correction for the variance of GEE estimators. Biometrical Journal, 45(4).

Neuhaus, J. M. and Kalbfleisch, J. D. (1998). Between- and Within-Cluster Covariate Effects in the Analysis of Clustered Data. Biometrics, 54(2).

Paul, S. and Zhang, X. (2014). Small sample GEE estimation of regression parameters for longitudinal data. Statistics in Medicine, 33(22).

Pedroza, C. and Truong, V. T. T. (2017). Estimating relative risks in multicenter studies with a small number of centers - which methods to use? A simulation study. Trials, 18(1).

Qaqish, B. F. (2003). A family of multivariate binary distributions for simulating correlated binary variables with specified marginal means and correlations. Biometrika, 90(2).

Rostrup, E., Gouw, A. A., Vrenken, H., Van Straaten, E. C., Ropele, S., Pantoni, L., Inzitari, D., Barkhof, F., and Waldemar, G. (2012). The spatial distribution of age-related white matter changes as a function of vascular risk factors-Results from the LADIS study. NeuroImage, 60(3).

Sachdev, P., Wen, W., Chen, X., and Brodaty, H. (2007). Progression of white matter hyperintensities in elderly individuals over 3 years. Neurology, 68(3).

Sharples, K. and Breslow, N. (1992). Regression analysis of correlated binary data: Some small sample results for the estimating equation approach. Journal of Statistical Computation and Simulation, 42(1-2).

Sherman, M. and Cessie, S. l. (1997). A comparison between bootstrap methods and generalized estimating equations for correlated outcomes in generalized linear models. Communications in Statistics Part B: Simulation and Computation, 26(3).

Veldsman, M., Kindalova, P., Husain, M., Kosmidis, I., and Nichols, T. E. (2020). Spatial distribution and cognitive impact of cerebrovascular risk-related white matter hyperintensities. NeuroImage: Clinical, 28.

Wang, Y. G. and Carey, V. (2003). Working correlation structure misspecification, estimation and covariate design: Implications for generalised estimating equations performance. Biometrika, 90(1).

Wardlaw, J. M., Smith, E. E., Biessels, G. J., Cordonnier, C., Fazekas, F., Frayne, R., Lindley, R. I., O’Brien, J. T., Barkhof, F., Benavente, O. R., Black, S. E., Brayne, C., Breteler, M., Chabriat, H., DeCarli, C., de Leeuw, F. E., Doubal, F., Duering, M., Fox, N. C., Greenberg, S., Hachinski, V., Kilimann, I., Mok, V., Oostenbrugge, R. v., Pantoni, L., Speck, O., Stephan, B. C., Teipel, S., Viswanathan, A., Werring, D., Chen, C., Smith, C., van Buchem, M., Norrving, B., Gorelick, P. B., and Dichgans, M. (2013). Neuroimaging standards for research into small vessel disease and its contribution to ageing and neurodegeneration. Lancet Neurology, 12(8).

Wardlaw, J. M., Valdés Hernández, M. C., and Muñoz-Maniega, S. (2015). What are white matter hyperintensities made of? Relevance to vascular cognitive impairment. Journal of the American Heart Association, 4(6).

Westgate, P. M. and Burchett, W. W. (2017). A Comparison of Correlation Structure Selection Penalties for Generalized Estimating Equations. American Statistician, 71(4).

Yelland, L. N., Salter, A. B., and Ryan, P. (2011). Performance of the modified poisson regression approach for estimating relative risks from clustered prospective data. American Journal of Epidemiology, 174(8).

Zhang, J. and Yu, K. F. (1998). What’s the relative risk? A method of correcting the odds ratio in cohort studies of common outcomes. Journal of the American Medical Association, 280(19).

Ziegler, A. and Vens, M. (2010). Generalized estimating equations: Notes on the choice of the working correlation matrix. Methods of Information in Medicine, 49(5).

Zou, G. (2004). A Modified Poisson Regression Approach to Prospective Studies with Binary Data. American Journal of Epidemiology, 159(7).

Zou, G. Y. and Donner, A. (2013). Extension of the modified Poisson regression model to prospective studies with correlated binary data. Statistical Methods in Medical Research, 22(6):661–670.

